# SPIKEPIPE: A metagenomic pipeline for the accurate quantification of eukaryotic species occurrences and intraspecific abundance change using DNA barcodes or mitogenomes

**DOI:** 10.1101/533737

**Authors:** Yinqiu Ji, Tea Huotari, Tomas Roslin, Niels Martin Schmidt, Jiaxin Wang, Douglas W. Yu, Otso Ovaskainen

**Author notes:** Contributed equally to the paper (Yinqiu Ji, Tea Huotari). **Statement of authorship:** NMS has been involved in running the BioBasis sampling program for more than twenty years. TR, NMS, DWY, and OO conceived the study and its design. TH led the work in generating all the DNA samples and YJ led the work in assembling and annotating the mitogenomes for the mitochondrial genome reference database. TH led the work in generating the mock communities and bulk samples, with contributions from YJ and JW. YJ and DWY developed the molecular and bioinformatic methods. OO led the modelling of the data. TR and OO wrote the first draa of the manuscript, and all authors contributed substantially to its further improvement. **Data accessibility statement:** Should the manuscript be accepted, the data supporting the results will be archived in an appropriate public repository (Dryad), and the data DOI will be included at the end of the article. The bioinformatic and R scripts and associated data tables will also be made available on github.com. **Name and complete mailing address of the persons to whom correspondence should be sent:** Prof. Otso Ovaskainen Organismal and Evolutionary Biology Research Programme, University of Helsinki, P.O. Box 65, FI-00014, Helsinki, Finland and Centre for Biodiversity Dynamics, Department of Biology, Norwegian University of Science and Technology, N-7491, Trondheim, Norway Prof. Douglas W. Yu School of Biological Sciences, University of East Anglia, Norwich Research Park, Norwich, Norfolk, NR47TJ UK and State Key Laboratory of Genetic Resources and Evolution, Kunming Institute of Zoology, Chinese Academy of Sciences, Kunming, Yunnan, 650223, China and Center for Excellence in Animal Evolution and Genetics, Chinese Academy of Sciences, Kunming Yunnan, 650223 China.

## Abstract

The accurate quantification of eukaryotic species abundances from bulk samples remains a key challenge for community ecology and environmental biomonitoring. We resolve this challenge by combining shotgun sequencing, mapping to reference DNA barcodes or to mitogenomes, and three correction factors: (1) a percent-coverage threshold to filter out false positives, (2) an internal-standard DNA spike-in to correct for stochasticity during sequencing, and (3) technical replicates to correct for stochasticity across sequencing runs. The SPIKEPIPE pipeline achieves a strikingly high accuracy of intraspecific abundance estimates (in terms of DNA mass) from samples of known composition (mapping to barcodes R^2^=0.93, mitogenomes R^2^=0.95) and a high repeatability across environmental-sample replicates (barcodes R^2^=0.94, mitogenomes R^2^=0.93). As proof of concept, we sequence arthropod samples from the High Arctic, systematically collected over 17 years, detecting changes in species richness, species-specific abundances, and phenology. SPIKEPIPE provides cost-efficient and reliable quantification of eukaryotic communities.

## 1. Introduction

The key dimensions of ecological community structure are species composition and species abundances (Vellend 2010). To survey communities of animals, plants, and fungi, researchers have traditionally relied on morphological characters. However, a technological revolution is beginning to replace this labor-intensive approach with spectral, acoustic, and molecular data (Bohan *et al*. 2017; Bush *et al*. 2017; Ovaskainen *et al*. 2018; Schulte to Bühne *et al*. 2018), offering replicable and efficient ways to estimate change in community structure over time, space, and environmental gradients.

We focus here on the use of DNA-sequence information to estimate eukaryotic species composition and abundance from mixed-species samples, such as bulk samples of invertebrates (Ji *et al*. 2013; Hering *et al*. 2018; Pawlowski *et al*. 2018), or water or air samples that have been filtered to capture environmental DNA (Bohmann *et al*. 2014; Abrego *et al*. 2018; Goldberg *et al*. 2018). In particular, we aim at quantifying within-species, spatiotemporal variation in a set of known target species, as measured by variation in the occurrence and mass of marker DNA sequences. This is the type of information typically sought in applied biomonitoring.

The two main approaches to identify eukaryotic species from mixed-species samples are known as ‘metabarcoding’ and ‘mitogenomics’ (Tang *et al*. 2015; Crampton-Platt *et al*. 2016; Bush *et al*. 2017; Bista *et al*. 2017). Metabarcoding uses PCR to amplify short, taxonomically informative ‘DNA barcode’ sequences from mixed-species samples. These amplicons are then sequenced, and the reads are assigned to taxonomies by matching to barcode reference databases. Metabarcoding is associated with low per-sample cost, because samples are individually tagged during PCR so that multiple samples can be pooled before being prepared for sequencing (‘library prep’) and because even a low sequencing depth of a few thousand reads is sufficient for characterizing the dominant species of a sample. This means that many samples can be included in the same sequencing run. Metabarcoding also has two other important advantages: access to well-populated reference databases, such as BOLD (Ratnasingham & Hebert 2007) and Midori (Machida *et al*. 2017), and the fact that PCR amplification allows the detection of species present only as low-concentration environmental DNA (eDNA), such as trace animal tissue in water. However, metabarcoding has two important limitations: susceptibility to sample contamination, due to PCR amplification of stray DNA (precisely what makes metabarcoding useful for eDNA in the first place), and loss of quantitative information, due to primer and polymerase biases (Yu *et al*. 2012; Piñol *et al*. 2015, 2018; Nichols *et al*. 2018; Deagle *et al*. 2018a; Lamb *et al*. 2019).

By contrast, mitogenomics is a variant of metagenomics and is based on the shotgun-sequencing of genomic DNA from a bulk sample, followed by mapping the reads onto a set of reference mitochondrial genomes, each of which serves as a ‘super barcode’ for a species (Crampton-Platt *et al*. 2016). Mitogenomics is more expensive than metabarcoding, because each sample must be individually library prepped, because samples must be sequenced more deeply than for metabarcoding, and because compiling a mitogenome reference database imposes additional costs for specimen acquisition, sequencing, and assembly. Mitogenomics is also unsuitable for eDNA because library prep requires sample DNA to be of high quality and quantity, and because mapping is only efficient if the sample consists mostly of the target organisms (but see Wilcox *et al*. 2018). However, mitogenomics has two key advantages over metabarcoding, due to not using PCR: mitogenomics is robust to sample contamination, because stray DNA is detected at low levels, if at all, and can thus be ignored. Also, mitogenomics preserves quantitative information, as the proportion of reads that map to a species’ mitogenome correlates with that species’ relative biomass in the sample (Zhou *et al*. 2013; Gómez-Rodríguez *et al*. 2015; Tang *et al*. 2015; Bista *et al*. 2017).

Thus, mitogenomics promises the ability to measure *both* of a community’s key dimensions, species composition and abundances. However, to date, reported correlations between read and abundance frequencies in samples of known composition have been noisy, and the shape of the relationship has varied idiosyncratically. For instance, analysing mixed-species communities, Zhou *et al*.’s (2013) pioneering study reported an R^2^ of 63% for a curvilinear regression, Tang *et al*. (2015) reported a linear-regression R^2^ of 25%, Gómez-Rodríguez *et al*. (2015) reported an R^2^ of 64% for a linear log-log regression, and Bista *et al*. (2017) returned R^2^ values between 45% and 87%, using logistic or linear models to fit different species.

Here we describe a step-change improvement in the mitogenomic pipeline, which performs well even when the mapping targets consist only of short DNA barcode sequences. To achieve accurate estimates of intraspecific variation in abundance, the pipeline employs a set of filters and internal standards: (1) a percent-coverage threshold to filter out false-positive mappings, (2) an internal-standard DNA spike-in to correct for sequencing-depth stochasticity across samples within a sequencing run, and (3) technical replicates to allow correction for sequencing-depth stochasticity across sequencing runs. We illustrate the pipeline by applying it to a time series (1997–2013) of piwall-trap samples from the high Arctic, the Zackenberg Valley in Northeast Greenland (Schmidt *et al*. 2016a; Christensen *et al*. 2017).

## 2. Materials and methods

SPIKEPIPE consists of five steps, which span from the wet lab to bioinformatics to statistical analysis (Box 1), and which are co-designed to maximize accuracy in species detection and to preserve abundance information. Step 1 consists of compiling a barcode or mitogenome reference database to be used as the mapping target. Step 2 consists of constructing, standardizing, and sequencing a set of mock communities to be used for calibration. Step 3 consists of standardizing and sequencing environmental samples (here, the Zackenberg samples). Step 4 consists of mapping the reads from the mock communities and environmental samples against the reference database and using these mappings to compute predictor values for species occurrence and abundance. Step 5 consists of using the mock data to fit statistical models that predict species occurrences and abundances, and applying these models to predictor values computed from the environmental data. We use DNA mass (ng of mitochondrial DNA of a species in the extracted sample) as our abundance measure, which is a proxy for species biomass. Below, we first introduce the Zackenberg case study, then detail Steps 1–5, and finally evaluate and compare the performance when using DNA barcodes versus mitogenomes to estimate community structure, both for the mock and the real environmental samples.

### Box 1.

An overview of the SPIKEPIPE metagenomic pipeline for quantifying eukaryotic species presences and abundances from mixed-species bulk samples. SPIKEPIPE maps shotgun-sequenced reads from the sample against whole or partial mitogenomes, including DNA barcode sequences (typically the 3’ portion of mtCOI for animals). Step 1 consists of compiling the barcode or mitogenome reference database (for barcodes, reference sequences can be downloaded from global databases such as BOLD; Ratnasingham & Hebert 2007). Step 2 consists of constructing, standardizing, and sequencing a set of mock communities to calibrate SPIKEPIPE. Step 3 consists of standardizing and sequencing a set of environmental bulk samples. Step 4 consists of mapping the reads from the mock communities and environmental samples against the reference database and using these mapping data to compute predictor values of species occurrence and abundance: *PC*, *FSL*, *SPECIES*, *RUN*. Step 5 uses the predictor values computed for the mock communities, for which we know the true species occurrences and abundances, Y(mock), to fit statistical models that predict species occurrences and abundances. These calibrated models are then applied to predictor values computed from the environmental mapping data to estimate species occurrences and abundances in the environmental samples, Y(environmental). Finally, the estimated species occurrences and abundances are used for modelling community change (Figs. 1, 2).

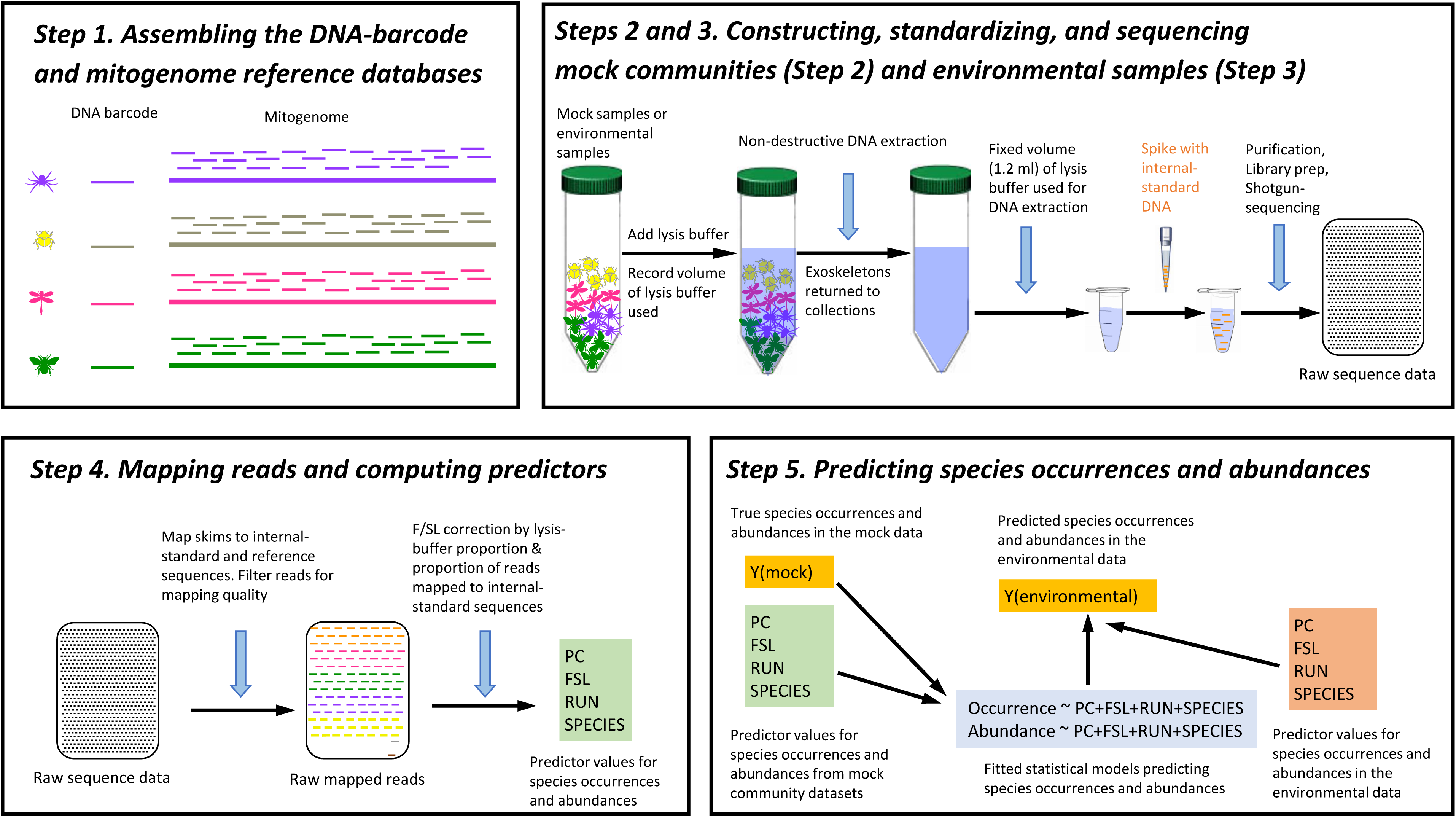

### Zackenberg Valley case study

The environmental dataset comes from a time series of yellow-piwall-trap samples collected as part of the BioBasis monitoring program at the Zackenberg Research Station (Schmidt *et al*. 2016a; Christensen *et al*. 2017), located in the High-Arctic zone of northeast Greenland (74°28’ N; 20°34’ W). Spiders and insects have been collected weekly throughout the summer from 1996 onwards from various tundra habitats. These samples have over the years been sorted to higher taxonomic rank (mostly family level), analyzed at this low taxonomic resolution (e.g. Høye *et al*. 2007, 2013, 2014; Reneerkens *et al*. 2016; Schmidt *et al*. 2016b, 2017), and warehoused as taxonomically sorted sub-samples in ethanol at room temperature in Museum of Natural History, Aarhus, Denmark (Supplement Text S3). The BioBasis protocols for sample collection, sorting, and storage have introduced cross-sample contamination (Supplement Text S3), making the samples unsuitable for metabarcoding. We use the subset of samples collected weekly in three yellow piwall traps in a mesic heath habitat, from 1997 to 2013 (Supplement Text S3), during which time summer has doubled in length (see Kankaanpää *et al*. 2018).

#### Step 1. Compiling the DNA-barcode and mitogenome reference databases

For the DNA-barcode database, we use the 406 insect and spider BIN sequences (Barcode Identification Numbers) compiled by Wirta *et al*. (2016) at Zackenberg (lengths range from 453 to 654 bp, mean 613). For the mitogenome database, we assembled 308 mitogenomes (lengths range from 5768 to 18625, mean 15720). The protocol for assembling mitogenomes is standard (Gómez-Rodríguez *et al*. 2015; Tang *et al*. 2015; Crampton-Platt *et al*. 2016; Bista *et al*. 2017), so we provide here a summary, with details in Supplement Text S1. Each voucher specimen was individually library-prepped and shotgun-sequenced to prevent chimeric assemblies and to preserve the option of using the nuclear-DNA reads in the future (cf. Sarmashghi *et al*. 2018). We successfully assembled 283 mitogenomes from these data, with a further 25 mitogenomes assembled out of a mixture of these data and the mixed-species piwall-trap samples’ data. Finally, we used COI from each mitogenome to confirm species identities, matching them to the BIN sequences of Wirta *et al*. (2016). Of the resulting 308 mitogenomes, 282 are in one contig, and the remaining 26 are in two or three contigs. 273 mitogenomes have all 13 protein-coding genes, and the remaining 35 have one or more missing or incomplete protein-coding genes (details in Supplement Table S1). 12S, 16S, and D-loop were seldom assembled, so these were omitted from all mitogenomes.

**Table 1.** True- and false-positive and negative observations of species occurrence in the mock-community experiment. The table shows the total numbers of species that were present (+) or absent (-), pooled over the 14 mock-communities (True cases). The next two rows report the number of inferred species presences based on (1) the lax criterion that ≥ 1 reads were mapped to a focal species and (2) the stringent criterion that mapping percent-coverage, PC, exceeded a threshold PC_0_. Note that even with the lax criterion, only 0.5% (barcode targets) or 3% (mitogenome targets) of truly absent species were identified falsely as present. With the stringent criterion, there were zero false positives.

|  | Barcode (+) | Barcode (-) | Mitogenome (+) | Mitogenome (-) |
| --- | --- | --- | --- | --- |
| True cases | 277 | 6219 | 277 | 4651 |
| $\geq 1$ read | 269 (97%) | 31 (0.5%) | 277 (100%) | 138 (3%) |
| $PC > PC_0$ | 264 (95%) | 0 (0%) | 269 (97%) | 0 (0%) |

#### Step 2. Constructing, standardizing, and sequencing mock communities

To calibrate SPIKEPIPE and estimate its accuracy, we constructed two kinds of mock communities using known input-DNA amounts of Zackenberg species: ‘mock-even’ and ‘mock-gradient’ communities (detailed in Supplement Table S2). We created six mock-even communities. For each, we used 20 species (19 Diptera, one spider), with equal amounts of DNA from each of the 20 species. This taxonomic composition reflects realistic sample compositions in the High Arctic, which is dominated by Diptera (Wirta et al. 2016). To mimic variation in sample absolute biomasses, two mock-evens used 50 ng DNA per species, two used 100 ng, and two used 200 ng (each creating a technical-replicate pair). We also created three mock-gradient communities. For each, we used 19 input species (all Diptera), with the first species represented by 20 ng of DNA, increasing geometrically by a factor of 1.3 until the most abundant species, at 2698.9 ng, a 135-fold range. The mock-gradient samples mimic the common situation where a sample contains both abundant and rare species (here, by DNA mass). We further included two negative controls without any species. The pooled set of mock communities allowed us to address the following questions: (1) *replicability*: do technical replicates return the same results?; (2) *within-species quantification*: for a given species, does the number of mapped reads correlate with the amount of input DNA?; (3) *across-species quantification*: do different species with the same input DNA return the same number of mapped reads?; and (4) *sensitivity*: can rare species be detected?

##### Internal-standard DNA

The fraction of the reads in a sample that maps to a species is an estimate of that species’ relative biomass in the sample, but to estimate that species’ absolute abundance (and allow comparisons across samples), we should correct for stochasticity in sequencing depth across samples. To do so, we used a fixed amount of an internal-standard DNA to ‘spike’ a 1.2 ml aliquot of lysis buffer from each of the mock samples (Box 1), so samples with more reads of the internal standard can be down-weighted (see Smets *et al*. 2016; Deagle *et al*. 2018b; Tkacz *et al*. 2018 for PCR-based protocols). To make the internal standard, we PCR-amplified and pooled fixed amounts of COI-barcode-amplicon DNA (658 bp) from three insect species collected in China and for which there are no confamilials in Greenland: 0.2 ng of *Bombyx mori* (Lepidoptera: Bombycidae), 0.4 ng of an unnamed beetle species (Coleoptera: Mordellidae), and 0.8 ng of another unnamed beetle species (Coleoptera: Elateridae) (For the PCR protocol, see Supplement Text S2.2). We used three species to check for error and degradation; the mapped reads from the three species should be found in the input ratio of 1:2:4. If, for instance, one of the species is not at the correct ratio with the other two, we can omit that species’ reads. Aaer spiking each mock sample, we extracted and purified the DNA and sent the mock samples for individual library prep and sequencing at the Earlham Institute (Supplement Text S2 for details). We sequenced the 6 mock-evens twice because one of the 50-ng mocks failed the first time, resulting in 11 total mock-even and 3 mock-gradient datasets, as well as the 2 negative controls.

**Table 2.**
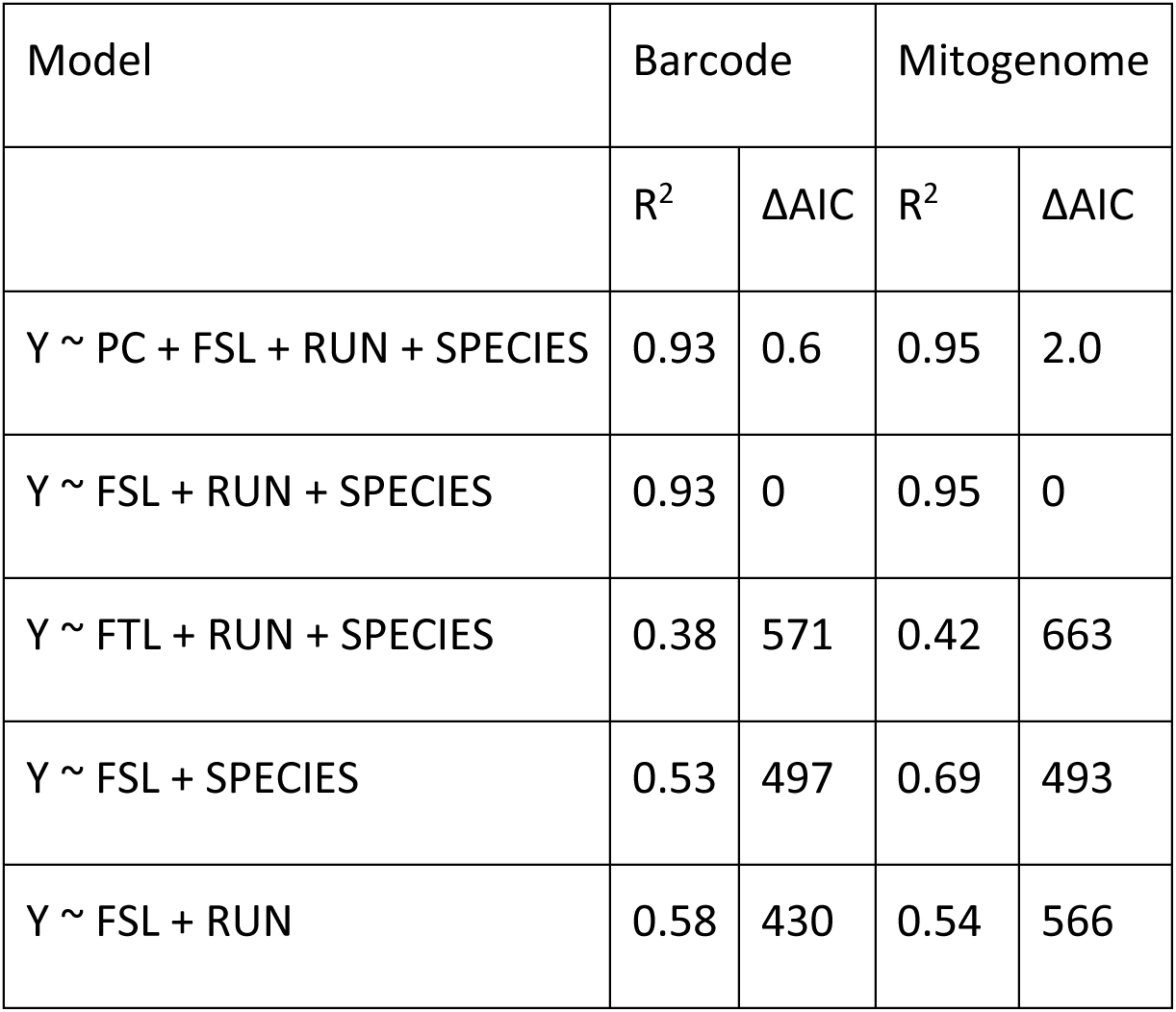
Variation in species abundance in the mock-community experiment explained by alternative models. In all models, the response variable Y is the log-transformed amount of DNA used for the focal species as input, and the columns show the proportion of explained variance R^2^ and ΔAIC (relative to the model with lowest AIC).

#### Step 3. Constructing, standardizing, and sequencing environmental samples

In 2016 and 2017, we non-destructively extracted DNA from the warehoused taxonomic subsets of 493 trap-week samples collected from 1997 to 2013 (except for 2010, the samples of which had been lost in transport from Greenland) (Supplement Text S4.1). We then pooled lysis-buffer solutions to reconstruct the 493 complete trap-week samples, aliquoted 1.2 ml from each sample, spiked each aliquot with 0.2 ng, 0.4 ng, and 0.8 ng of the three internal-standard species (from Step 2), extracted and purified the DNA, and sent them for individual library prep and sequencing at the Earlham Institute. The samples were spread over three sequencing runs, so we included 4 or 5 technical replicates per year of samples from the previous two runs in the final run, to allow correction for stochasticity in sequencing depth across runs (Supplement Text S4.2).

In a variant protocol, for years 1997-1999 and 2011-2013, the first samples that we sequenced, we used 10 ng, 20 ng, and 40 ng of the three internal-standard species. We subsequently determined that the internal-standard DNA represented a too-large proportion of the resulting reads (mean 10.9%), so we library-prepped and sequenced the remaining DNA from the aliquots a second time to increase data. Altogether, we successfully sequenced 728 samples: 14 mocks, 712 trap-week samples (original and technical-repeat aliquots), and two negative controls (which returned only the internal-standard species) (Supplement Table S4.2.1).

#### Step 4. Mapping reads and computing predictors

All sequencing outputs (mock communities, negative controls, and the Zackenberg samples) were processed as follows (details in Supplement Text S5). We removed Illumina adapter sequences using *TrimGalore* 0.4.5 (Martin 2011) and used *minimap2* 2.10 in short-read mode (Li 2018) to map the paired-end reads against either the DNA-barcode or the mitogenome reference database, plus the three internal-standard sequences. We used *samtools* 1.5 (Li *et al*. 2009) to exclude reads that mapped as secondary or supplementary alignments and to include only paired-end reads mapped at quality ≥48, in the correct orientation, and at approximately the correct distance apart (‘proper pairs’).

*PC, FSL, RUN, SPECIES* – If a species is truly present in a sample, we expect reads from that sample *to map along the length* of that species’ DNA-barcode or mitogenome, not just to one segment, resulting in a higher percentage coverage (henceforth *PC*) (Supplement Figure S5.1). We used *bedtools* 2.27.1 (Quinlan & Hall 2010) to calculate PC as the fraction of positions covered by one or more mapped reads. We expected the fraction of reads from a sample that map to a species’ barcode or mitogenome to be correlated with that species’ relative abundance. However, the absolute number of mapped reads in a sample is determined by that sample’s dataset size, which is affected by numerous random factors from DNA extraction to sequencing. We used an internal standard with the aim of correcting for these random variations and recovering each sample’s original biomass, allowing calculation of each species’ absolute abundance per sample (Box 1). To do so, we added the same amount of internal-standard DNA to the fixed aliquot of each sample before DNA extraction (see *Internal-standard DNA*), and kept track of the fraction of total lysis buffer that the aliquot represents. We defined the quantity *FSL* for each species in each sample, defined as log(*F/SL*), where *F* is the number of reads mapped to a focal species (unit: sequence count), *S* is the “spike” (unit: sequence count/DNA mass), and ( is the fraction of lysis buffer represented by the aliquot (unitless). The spike *S* was first computed for each of the three internal standards as number of reads mapped to the internal standard divided by the input amount of DNA, and then averaged over the three internal standards. To test the importance of the internal standard, we also computed *FTL*, defined as log(*F/SL*), where *F* and *L* are the same as above, and *T* is the total number of sequences in a sample’s dataset. As the mapping from *PC* and *FSL* to species presences and abundances might also vary among sequencing runs, we used the identity of the three sequencing runs (henceforth RUN), as an additional predictor, implemented as a factor. Finally, as sequencing success could be idiosyncratically species-specific, e.g. species varying in their mitochondrial-to-nuclear-DNA ratios or in how much tissue is released into solution during lysis, we considered the identity of the species (henceforth *SPECIES*) as an additional predictor, implemented as a factor. To summarize, the outcome of Step 4 is two datasets of values for *PC*, *FSL*, *RUN,* and *SPECIES*, computed aaer mapping reads from the pooled mock-community dataset against the DNA-barcode and the mitogenome reference databases.

#### Step 5. Predicting species occurrences and abundances

In this step, statistical models that predict species occurrences and abundances were calibrated with the mock-sample datasets and then applied to the environmental data. First, we used logistic regression to model species presence/absence as a function of *PC*, *FSL*, *RUN*, and *SPECIES*, of which we expected *PC* to be the most important predictor. Second, we used a linear regression with the same predictors to model variation in abundance conditional on presence (the log-transformed amount of DNA included in the mock community), with the expectation of *FSL* being the most important predictor. Note that we used log-transforms in our models because the amounts of DNA in both the mock samples and the environmental samples can vary over orders of magnitude. In such a case, the researcher is likely to be interested in quantifying proportional differences among samples, e.g. “sample A has 100 times more and sample B has 10 times more DNA of the focal species than sample C has”. Moreover, we log-transformed both the response variable and the main predictor (FSL) to make them compatible with each other. Thus, the theoretical expectation for the slope value of *FSL* is 1.0, because that corresponds to the number of (SL-corrected) mapped reads being directly proportional to the amount of input DNA.

To examine the information gained from the addition of the spike, we replaced in the best model *FSL* by *FTL*. To examine how much accuracy is lost if the results are not calibrated per run, we dropped *RUN* from the model. To examine how well the abundance estimates are calibrated across species, we examined performance of a model variant where *SPECIES* was dropped. Further, to examine if species effects can be explained by their mitogenome lengths, we regressed the estimated effects of *SPECIES* from the abundance model on log-transformed mitogenome length, reasoning that longer mitogenomes should attract more reads. Finally, the models calibrated with mock community data were used to predict species occurrences and abundances in the environmental data (Box 1).

### Evaluating SPIKEPIPE’s replicability with re-sequenced samples

After performing steps 1–5 of SPIKEPIPE described above, we tested its replicability by comparing results between technical replicates. If a sample was sequenced more than twice (see Table S4.2.1), we considered all pairs of replicates, so that a sample sequenced in runs X, Y and Z contributed to the comparisons (X,Y), (X,Z) and (Y,Z). To evaluate the robustness of detecting species presence, we counted how oaen a species was inferred to be present or absent in both runs of comparison (agreement) or present in only one (disagreement). To evaluate the robustness of species abundance estimation, we used only cases where a species was classified as present in both runs of comparison. For each pair of runs, we used a linear model to estimate how well the abundance estimates obtained from one run explained variance in abundance estimates in the second run. We performed this analysis either controlling or not controlling for the effect of the specific run, to examine whether run-specific correction factors are needed.

### Evaluating SPIKEPIPE’s utility for making ecological inference

Finally, we tested SPIKEPIPE’s power to detect ecological change, and how this power differs between mapping against reference DNA barcodes or reference mitogenomes. To this aim, we merged the results from the four runs by assuming that a species is present in a sample if it was inferred to be present in any of the runs containing that sample, and we used the average of run-corrected abundance estimates as the final estimate for abundance. We then normalized abundance within each species across all samples to mean zero and unit variance, so that the unit of the abundance is within-species deviation from the species’ mean. We summed inferred presences over all species to estimate species richness per trap-week, and we averaged normalized abundances over all species to estimate community-wide variation in abundance. To test whether these richness and abundance data convey signals of change in a warming Arctic, we used Poisson regression to model species richness per trap-week, and linear regression to model mean abundance conditional on presence. In both cases, the data points (n=493) are weekly samples from the three traps in the years 1997–2013. We used as the candidate predictors the continuous effect of the year (to capture a linear trend), the Julian date and its square (to capture phenological variation within year), the interaction between year and Julian date (to capture change in phenological timing), and the trap (to capture technical variation among the three traps). We performed variable selection in all models using the Akaike Information Criterion (AIC).

## 3. Results

### 3.1. Mock community analyses: species composition and abundances

Mapping to short barcodes and long mitogenomes were both highly successful in identifying species occurrences in the mock samples, even though these included the mock-gradients with their rare species: the barcode approach detected 97%, and the mitogenome approach detected 100% of the species that were actually present (Table 1). The false-positive rates were also very low even when using the lax criterion of lettng just one mapped read infer species presence (Table 1). Due to this high separation, fittng logistic regression models was not meaningful. Instead, we defined conservative thresholds *PC_0_* for percentage cover (set to *PC_0_*=0.5 for barcodes and *PC_0_*=0.1 for mitogenomes), so that applying the threshold condition *PC* > *PC_0_* resulted in zero false positives and thus gave strong evidence for species occurrence in a sample (Table 1, see also Tang *et al*. 2015). Sensitivity remained high when applying the criterion *PC* > *PC_0_*, with the barcode approach detecting 95% and the mitogenomics approach 97% of the species that were actually present.

In terms of sensitivity, we found a taxonomic difference. Among the two classes represented in the mock communities (19 Diptera, one spider), the spider performed worse than the other taxa in our analyses: in the mock data, the spider was present in 11 samples and absent in 5 samples. For the barcode approach, the presence-absence model predicted that the species was present only once, resulting in 1 true positive, 5 true negatives, and 10 false negatives. For the mitogenomics approach, the number of true positives was 3, and thus the number of false negatives was 8. This implies that further tests with species outside Diptera will be needed to validate the generality of our results.

Turning next to species abundances, we focused on the species inferred to be present by *PC* > *PC_0_*, attempting predict how abundant each was. Model selection with AIC supported keeping *FSL*, *RUN*, and *SPECIES* in the final model when mapping against either barcodes or mitogenomes. With these models, R^2^ equaled 0.93 for barcodes and 0.95 for mitogenomes (Table 2, Box 1), and the slope of *FSL* was 0.90 for both barcodes and mitogenomes, i.e. close to the theoretical expectation of 1.0. Models without the internal standard (i.e. using *FTL* rather than *FSL* as the predictor) performed poorly, as did models from which *RUN* or *SPECIES* was dropped (Table 2, Box 1). Variation among *SPECIES* effects in the final model was not explained by length of barcode (p=0.96) nor mitogenome (p=0.21). In total, these results imply that calibration between sequencing runs is necessary and that only *within*-species variation in abundance can be quantified accurately (i.e. ‘Species A is more abundant in this sample than in another’). To estimate *among*-species variation (i.e. ‘Species A is more abundant than species B’), it will be necessary to run mock communities with all target species of interest to estimate species-specific calibration factors.

### 3.2. Zackenberg analyses: replicability

Data based on mitogenomes provided a somewhat higher success rates than did data based on barcodes, as judged by two criteria: a higher number of species presences inferred, and a lower proportion of cases where a species was inferred to be present in one run only (Table 3). But overall, the numbers were surprisingly similar (Table 3), given that mean barcode length was only 3.9% of mean mitogenome length. The comparisons also suggest that the error rate in identifying species presence is somewhat higher in the environmental data (Table 3) than with the mock communities (Table 1), as may be expected given extra complexities with environmental data. The consistency of the run-corrected abundance estimates was almost as high as with the mock data: the abundance estimate from one run explained 94% (for barcodes) or 93% (for mitogenomes) of the variation of the abundance estimate derived from the other run (Fig. 1AD). Species of different taxonomic affinity varied slightly in performance (see Supplement S4.3). The consistency among replicate samples was somewhat higher for Araneae than for Diptera, and somewhat lower for Hymenoptera and Lepidoptera than for Diptera, but these latter two orders were generally too rare to make robust conclusions.

**Table 3.**
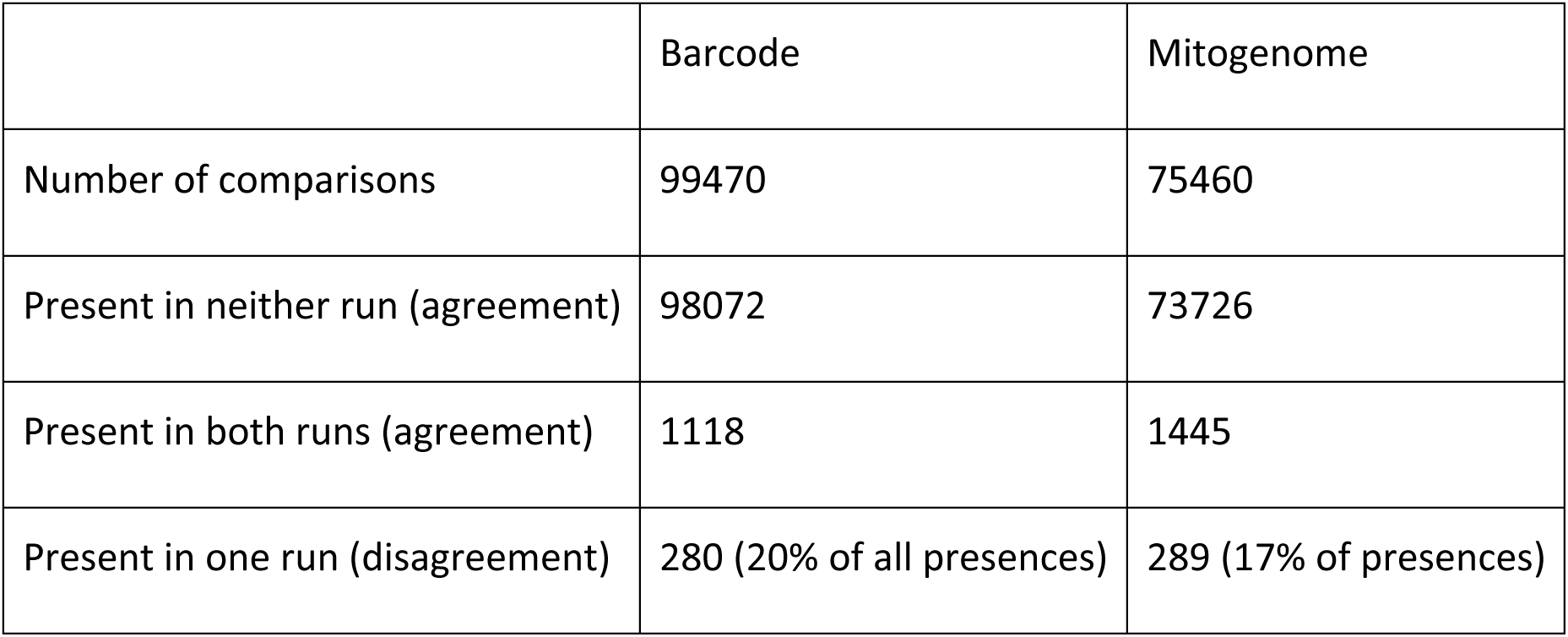
Consistency of re-sequenced environmental data. The table shows the consistency in species presences and absences between two sequencing runs, with the total number of comparisons equaling the total number of sample-by-species combinations. To evaluate the robustness of detecting species presence, we counted how often a species was inferred to be present or absent in both cases (agreement) or present in one case only (disagreement). The proportion of presences is counted as the fraction of all samples where the species was inferred to be present in at least one run.

### 3.3. Zackenberg analyses: ecological inference

Of the 371 species that were present in our DNA-barcode reference database and that were systematically sampled over the years, we observed across all samples 145 species at least once, and 72 species at least five times. Of the 308 species in our mitogenome reference database, we observed 148 species at least once, and 81 species at least five times. Even though our analyses with mitogenome reference database contained 17% fewer species, mapping to mitogenomes consistently revealed higher species richness (53 species observed on average per year) than did mapping to DNA barcodes (46 species observed on average per year; Fig. 2).

**Figure 1.**
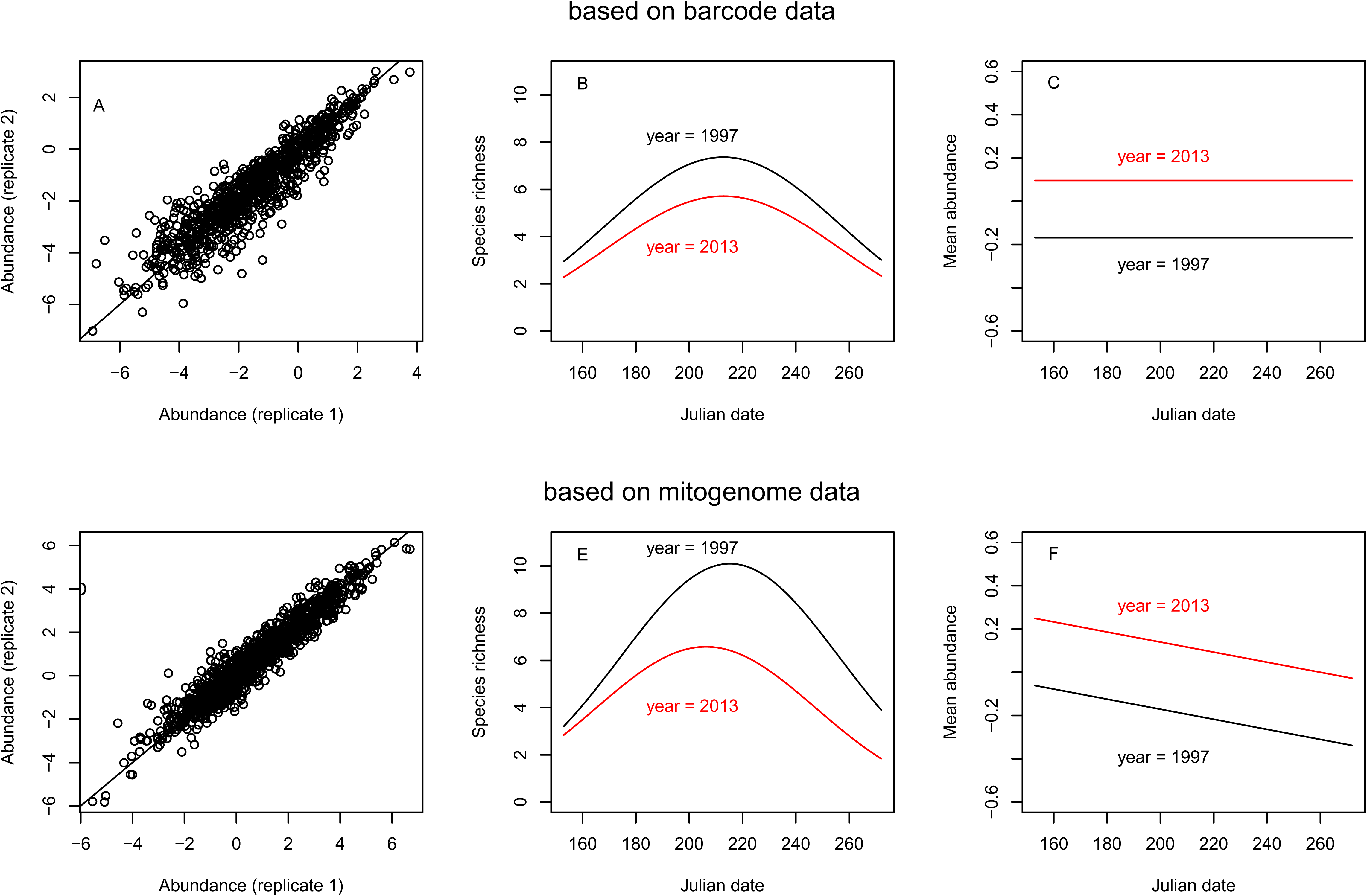
Results from the environmental Zackenberg arthropod samples from 1997 and 2013, collected with yellow piball traps and processed using the SPIKEPIPE pipeline. The upper panels (A-C) are based on mapping against DNA barcodes, and the lower panels (D-F) on mapping against mitogenomes. Panels A, D examine the technical validity of the data by comparing run-corrected abundance esdmates for species sequenced in technical replicates across sequencing runs. Panels B, E show the species richness per trap-week in 1997 (black line) and 2013 (red line), as predicted by a Poisson regression model fifed to weekly data on all years 1997–2013, as described in the main text. (For clarity, only the stardng and end years are shown in the figure.) Panels C, F show the mean abundances per week (in the unit of log-transformed DNA amount), normalized to zero mean and unit variance within each species) of those species that were present in 1997 and in 2013, as predicted by linear regression models described in the main text.

**Figure 2.**
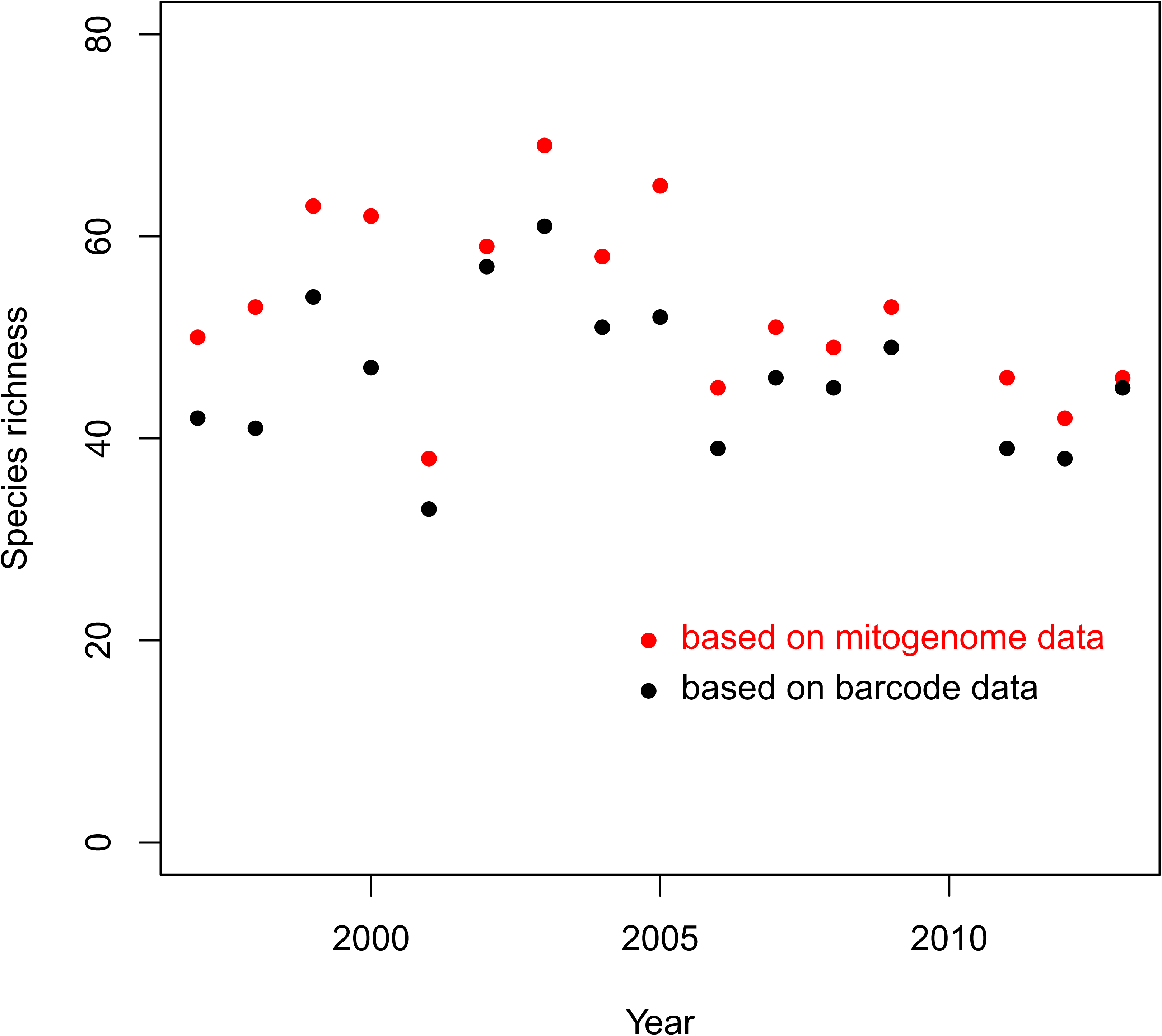
Total number of species detected per year in the Zackenberg samples, based on mapping to either DNA barcodes (black dots) or to mitogenomes (red dots).

We then contrasted the effects of using barcodes versus mitogenomes on estimates of community change, including only species that were observed at least five times. In our models of species richness *per trap-week*, we observed a peak in the middle of the summer and a decrease from 1997 to 2013 (Fig. 1BE), regardless of whether we used barcodes or mitogenomes. With mitogenomes (but not with barcodes), a statistical interaction between year and Julian season was retained, suggesting that peak species richness occurred earlier in 2013 than in 1997 (Fig. 1E). Both the barcode and the mitogenome datasets recorded greater total species abundance in 2013 over 1997, but only the mitogenome approach detected that community abundance decreased over the summer (Fig. 1CF).

### 4. Discussion

The harnessing of high-throughput DNA sequencing to infer species composition and species abundances is a huge opportunity for community ecology and environmental monitoring (Tang *et al*. 2015; Bush *et al*. 2017; Bista *et al*. 2017; Porter & Hajibabaei 2018). We report a step-change improvement in the accuracy and precision of the mitogenomic approach, achieved by combining shotgun sequencing with a percent-coverage threshold (*PC_0_*), an internal-standard DNA spike-in (*FSL*), and correction factors among sequencing runs (*RUN*; Box 1). The highest accuracy and precision are achieved with longer mitogenome targets but remain surprisingly high with short DNA-barcode targets (Table 1, 3; Box 1, Fig. 1), and both approaches recover signals of structural change from real samples of arthropods at a high-Arctic site with marked environmental change (Figs. 1, 2).

### 4.1. High accuracy and precision

Our pipeline accurately inferred the presence and absence of species in mock samples. By selecting a conservative threshold for percentage cover (Table 1), we could completely eliminate false positives, while keeping false-negative rates only at 3% and 5% when mapping to mitogenomes and DNA barcodes, respectively. However, differences in the performance of Diptera and Araneae suggest that further tests will be needed to validate the generality of our results across a wider range of taxa. We could also accurately estimate within-species variation in abundance, achieving R^2^ values of 0.95 for mitogenomes and 0.93 for COI barcodes, and reaching almost direct proportionality between the number of mapped reads and species abundance (Fig. 1). These results are a major improvement over many previous studies (including our own), which have reported positive but noisy and idiosyncratic correlations between read number and biomass (Tang *et al*. 2015, Zhou *et al*. 2013; Gómez-Rodríguez *et al*. 2015; Bista *et al*. 2017). Overall, they seem to deliver on the key promise of mitogenomic approaches, adding quantitative information beyond that achieved by metabarcoding approaches (see Introduction).

We used as a unit of abundance DNA mass – i.e. nanograms of mitochondrial DNA of each species in a compound sample of DNA. Importantly, this is an absolute rather than a relative metric, allowing one to assess variation in the occurrence and concentration of DNA of the target species among samples, and thus to estimate within-species population changes. By measuring such changes for multiple species, one can detect quantitative changes in community structure. If desired, the unit of DNA mass can be converted into an estimate of the more traditional metric of individual counts per sample, given the availability of an estimate of mitochondrial DNA mass per individual per species.

Like any survey method, the method presented here will likely miss some species that are actually present in an environmental sample (Table 3), especially those present at low abundance. Thus, exhaustive sampling will require several sequencing runs, or at least a very high sequencing depth in a single run. When multiple sequencing runs are used, our results suggest that variation between runs can be almost completely accounted for, if including some samples in all runs to enable calibration. However, our tests are not exhaustive, and that this conclusion should thus be taken with caution. We also note that our mock samples were designed to reflect the High-Arctic fauna, which is dominated by the order, Diptera, and we have shown that it is possible to differentiate congener species in this order, which is an important requirement for understanding changes in this community. However, we caution that given the enormous diversity of the Arthropoda, our tests can only be considered partial, and we robustness should continue to be tested in other studies.

### 4.2 Mitogenomes versus DNA barcodes

We consistently detected more species when mapping to mitogenomes than to barcodes – even though we had 406 barcodes but only 308 mitogenomes (Table 1, Fig. 2). This difference was caused by the higher PC threshold needed to avoid false positives for barcodes (*PC_0_* = 0.5) than for mitogenomes (*PC_0_* = 0.1). Perhaps as a result, mitogenome datasets also appear to be more powerful for detecting change in community structure (Fig. 1EF).

However, the higher statistical power of a mitogenome reference dataset trades off against the higher cost of compiling it, especially if suitable barcode sequences are already available (Ratnasingham & Hebert 2007; Machida *et al*. 2018; Nilsson *et al*. 2018). A mitogenome reference dataset is likely justified if the aim is to repeatedly monitor a fixed species list, in which case the initial cost of assembling a mitogenome dataset can be amortized. Conversely, in species-rich areas, DNA barcodes might be the only feasible option, in which case our results suggest that it will be helpful to increase sequencing depth per sample, and possibly also the number of samples per location. For example, the new Illumina NovaSeq delivers 1.5–5 times the output of the Illumina 2500 that we used (Bleidorn 2017). Our per-sample sequencing cost on the Illumina 2500 ranged from 130€ to 220€ per sample, plus the LITE library cost of 5.6€ per sample. A high sequencing depth was required because in our samples, only ∼0.2-0.4% of the reads mapped to our 308 mitogenomes, the remainder representing the nuclear genomes or microbes, not counting the internal-standards. Baits could potentially help (Liu *et al*. 2015, Wilcox *et al*. 2018), but baits do introduce bias, and the enrichment factor will vary across samples, obscuring abundance information. A final consideration is that a barcode reference dataset is currently more likely to allow detection of unexpected species, since barcode datasets can more easily include species from outside the study area. However, it could eventually become routine to carry out high-volume *de novo* mitogenome assembly from bulk samples themselves (Crampton-Platt *et al*. 2015), allowing species discovery and detection from the same samples. A related strategy is targeted assembly of barcodes from shotgun-sequenced samples (Greenfield et al. 2019).

### 4.3. Changes in community structure correlated with Arctic warming

SPIKEPIPE revealed major changes in mean phenology and abundance in the Zackenberg arthropod community. While we will report detailed analyses elsewhere, the patterns identified in Fig. 1BCEF expand upon two morphology-based studies in Zackenberg. Loboda *et al*. (2018) individually identified 18,385 Muscidae flies into 16 species and found a decline in species diversity between 1996 and 2014 – but no statistically significant change in species abundances in the mesic heath habitat (the same habitat as our environmental samples). Bowden *et al*. (2018) individually identified 28,566 spiders into nine species and found a decline in some species but no change in others. Such differential responses across species are altering the structure of Arctic communities (Høye *et al*. 2014; Kankaanpää *et al*. 2018, Koltz *et al*. 2018), and the changes are percolating to functional associations, e.g. between plants and their pollinators (Schmidt *et al*. 2016b; Cirtwill *et al*. 2018). We clearly need species-resolution time series from full communities in order to understand the effects of climate change on biodiversity. An expansion of the morphological approach is infeasible, but our improvements in quantification, coupled with recent and imminent gains in cost-efficiency, now make the mitogenomic approach an attractive option for use in community ecology and in applied biomonitoring.

## Acknowledgments

Arthropod samples were provided by the Greenland Ecosystem Monitoring Programme. The authors gratefully acknowledge CSC *–* IT Center for Science Finland and the HPC team at UEA UK, for computational resources. We thank the Danish National High-Throughput DNA Sequencing Centre and the Earlham Institute for sequencing. We thank Jaana Kekkonen and Isabella Palorinne for help during the laboratory work. We received funding from the Academy of Finland (grants 276909 and 285803 to TR, grants 284601 and 309571 to OO), the Jane and Aatos Erkko Foundation Grant (OO and TR), the Research Council of Norway through its Centres of Excellence Funding Scheme (223257) to OO via Centre for Biodiversity Dynamics. We are indebted to the Danish Environmental Protection Agency for funding BioBasis Zackenberg over the years, and to the many field and lab assistants over the years. D.W. Yu and Y.Q. Ji were supported by the National Natural Science Foundation of China (41661144002, 31670536, 31400470, 31500305), the Key Research Program of Frontier Sciences, CAS (QYZDY-SSW-SMC024), the Bureau of International Cooperation (GJHZ1754), the Strategic Priority Research Program of the Chinese Academy of Sciences (XDA20050202, XDB31000000), the Ministry of Science and Technology of China (2012FY110800), the State Key Laboratory of Genetic Resources and Evolution (GREKF18-04) at the Kunming Institute of Zoology, the University of East Anglia, and the University of Chinese Academy of Sciences. The funders had no role in study design, data collection and analysis, decision to publish, or preparation of the manuscript.

## Supporting Information

Supporting Information includes additional information on constructing the mitochondrial genome reference database (Text S1), constructing, standardizing, and sequencing mock communities (Text S2), the environmental samples (Text S3), processing of bulk samples from the Zackenberg collection (Text S4), mapping reads against references (Text S5), and instructions on how to download the example data and run Step 4 (bioinformatics) of the pipeline (Text S6), and how to run Step 5 (statistics) of the pipeline (Text S7).

## Supplementary material to

### Text S1. Constructing the mitochondrial genome reference database

Like all metagenomic approaches, SPIKEPIPE relies on the availability of a comprehensive reference library. Only species represented in the reference library can be detected, since sample sequences are mapped against only the sequences present in the reference library. An obvious challenge therefore resides in acquiring tissue for a locally representative set of species. In the present case, this challenge was resolved by our long-term work on clarifying the full fauna of the target region (Wirta *et al*. 2015), allowing us to construct a comprehensive collection of identified voucher specimens and – for a majority of species – to obtain voucher extracts of DNA.

#### S.1.1. DNA extraction

In previous work, voucher samples were sent for Sanger barcoding at the Canadian Center of DNA Barcoding, University of Guelph (full details in Wirta *et al*. 2015). Here, sequences of the standard barcoding region of COI were clustered into 410 BINs (Ratnasingham & Hebert 2013). After double-checking the alignments of COI sequences manually, four BINs were removed because of gaps obviously resulting from sequencing errors, leaving 406 BINs. Of these, we did not attempt mitogenome assembly with Collembola (16 species) and Acari (19 species), since these taxa are tiny and thus unlikely to provide enough DNA for individual genome sequencing. In addition, they have been inconsistently stored from the Zackenberg samples (see S3.1 for more details).

Drawing on the voucher collection established by Wirta *et al*. (2015), we re-extracted genomic DNA from each of the 364 available species using the DNeasy Blood and Tissue Kit (Qiagen, Hilden, Germany). We applied a non-destructive DNA extraction protocol to the voucher specimens: specimens are soaked in the kit’s extraction buffer, and DNA is then purified from the soaking solution according to the protocol of the extraction kit. All the specimens used for constructing the mitogenome reference library are stored at the Department of Agricultural Sciences at the University of Helsinki.

#### S.1.2. Sequencing

Genomic DNA of 171 species was sent for shotgun sequencing at the Danish National High-throughput Sequencing Center (Copenhagen, Denmark), where TruSeq libraries with 500 bp insert size were built on the NeoPrep Library Prep System and sequenced at 125 bp PE on an Illumina HiSeq 2500. The remaining 193 species, which could not reach the minimum requirements for TruSeq library construction, plus 17 species for which sequencing at Copenhagen failed, were sent to Earlham Institute in Norwich, UK. Here, Low Input Transposase-Enabled (LITE) libraries with 400 bp insert size were built and sequenced at 250 bp PE on an Illumina HiSeq 2500.

#### S.1.3. Bioinformatics

The raw reads were corrected with *bfc* (-s 3g) (Li 2015). *TrimGalore* 0.4.4 (--length 50 --trim-n) (https://github.com/FelixKrueger/TrimGalore) (Martin 2011) was used to trim low-quality base calls from the 3’ end, to remove adapter sequences, and to omit short reads with a length of less than 50 bp. Most mitogenomes were assembled successfully by two assemblers: *IDBA-UD* (-r for 125 bp and –l for 250 bp) (Peng *et al*. 2012) and *SPADES* 3.11.0 (--only-assembler –meta -k 21,33,55) (Nurk *et al*. 2013). For species for which complete mitogenomes could not be created by *IDBA* and *SPADES*, several other assemblers were tried, including *SOAPdenovo*-Trans 1.03 (-K 31 –L 100 –t 1) (Xie *et al*. 2014), *SADBG* composed of *SparseAssembler* 2 (LD 0 k 31 g 15 NodeCovTh 2 EdgeCovTh 1 GS 16000) (Ye *et al*. 2012) and *DBG2OLC* v1 (k 31 KmerCovTh 0 MinOverlap 30 PathCovTh 3 LD 0) (Ye *et al*. 2016), *NOVOPlasty* 2.6.3 (K-mer = 39) (Dierckxsens *et al*. 2016) and *MITObim* 1.9 (-end 100 –quick --denovo) (Hahn *et al*. 2013). The resulting scaffolds were used as query by BLAST (-evalue 1e-7) against an Arthropoda mtDNA database downloaded from Genbank, and only those with a hit and a length no less than 3 kb were retained for subsequent analyses. In order to improve the quality of our mitogenome reference database, the filtered scaffolds were error-corrected with *pilon* 1.18 (Walker *et al*. 2014) by using the post-QC reads mapped with *bwa* (Li 2013). The COI sequences of Wirta *et al*. (2015) as included in the barcode reference library (above) were used as a reference to find the corresponding mitogenomes from the corrected filtered scaffolds with the function “Map to Reference” in GENEIOUS 11.0.4 (http://www.geneious.com). MITOS 2 (http://mitos2.bioinf.uni-leipzig.de) (Bernt *et al*. 2013) was used to annotate the mitogenomes. For species for which the resulting mitogenomes did not have all the 13 protein coding genes, the scaffolds obtained by all the assemblers were assembled using GENEIOUS assembler to get the longest mitoscaffolds. The protein-coding genes were extracted from the mitogenomes and translated to ensure correct translation frames in GENEIOUS. In order to get more mitogenomes, the sequence data of the bulk samples (see below) were also mined. The assembly pipeline is the same as the above, except that the correction step of *pilon*-*BWA* was skipped because *BWA* would have produced mapping hits from multiple species. Five mitogenomes were successfully assembled through this latter approach. In total, we assembled partial or full mitogenomes (considering protein-coding genes only) for 308 of the 371 Zackenberg insect and spider BINs.

**Table S1.**
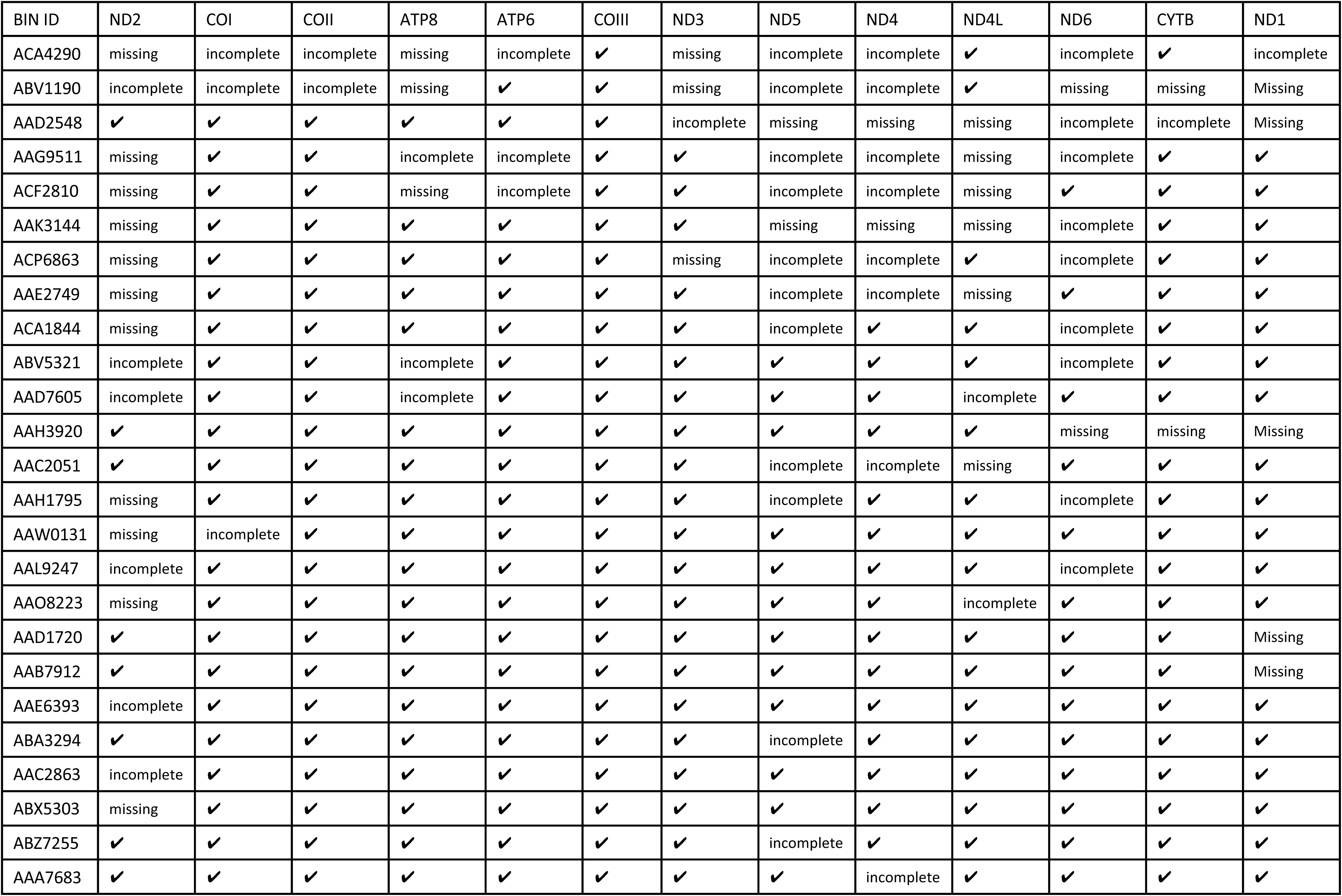
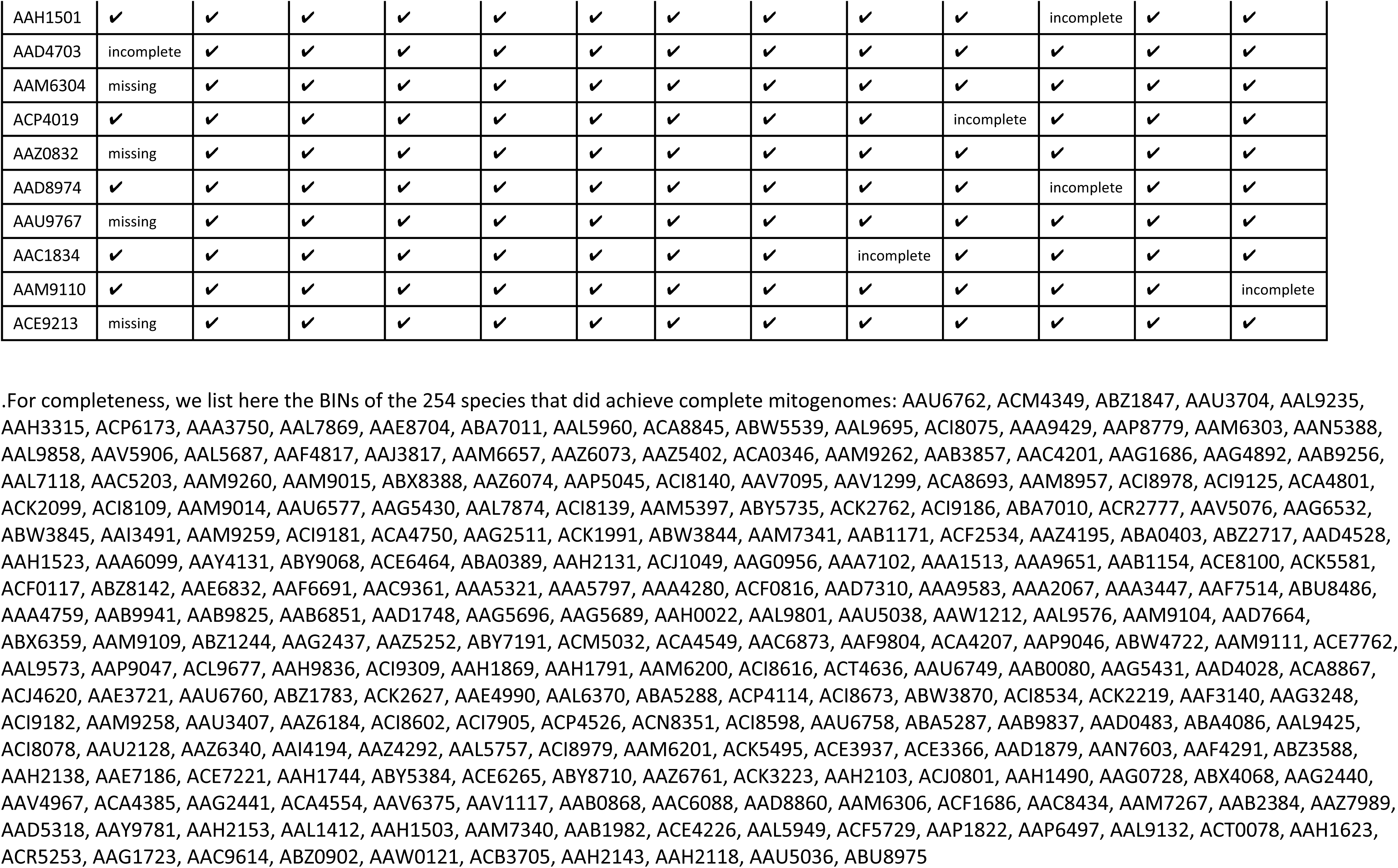
Sequence availability from 13 mitochondrial protein genes in 308 species from Zackenberg, Northeast Greenland. The 35 species that did not achieve complete mitogenomes, listed under their BIN ID numbers, with full taxonomies available in Wirta *et al*. 2015.

| BIN ID | ND2 | COI | COII | ATP8 | ATP6 | COIII | ND3 | ND5 | ND4 | ND4L | ND6 | CYTB | ND1 |
| --- | --- | --- | --- | --- | --- | --- | --- | --- | --- | --- | --- | --- | --- |
| ACA4290 | missing | incomplete | incomplete | missing | incomplete | ✓ | missing | incomplete | incomplete | ✓ | incomplete | ✓ | incomplete |
| ABV1190 | incomplete | incomplete | incomplete | missing | ✓ | ✓ | missing | incomplete | incomplete | ✓ | missing | missing | Missing |
| AAD2548 | ✓ | ✓ | ✓ | ✓ | ✓ | ✓ | incomplete | missing | missing | missing | incomplete | incomplete | Missing |
| AAG9511 | missing | ✓ | ✓ | incomplete | incomplete | ✓ | ✓ | incomplete | incomplete | missing | incomplete | ✓ | ✓ |
| ACF2810 | missing | ✓ | ✓ | missing | incomplete | ✓ | ✓ | incomplete | incomplete | missing | ✓ | ✓ | ✓ |
| AAK3144 | missing | ✓ | ✓ | ✓ | ✓ | ✓ | ✓ | missing | missing | missing | incomplete | ✓ | ✓ |
| ACP6863 | missing | ✓ | ✓ | ✓ | ✓ | ✓ | missing | incomplete | incomplete | ✓ | incomplete | ✓ | ✓ |
| AAE2749 | missing | ✓ | ✓ | ✓ | ✓ | ✓ | ✓ | incomplete | incomplete | missing | ✓ | ✓ | ✓ |
| ACA1844 | missing | ✓ | ✓ | ✓ | ✓ | ✓ | ✓ | incomplete | ✓ | ✓ | incomplete | ✓ | ✓ |
| ABV5321 | incomplete | ✓ | ✓ | incomplete | ✓ | ✓ | ✓ | ✓ | ✓ | ✓ | incomplete | ✓ | ✓ |
| AAD7605 | incomplete | ✓ | ✓ | incomplete | ✓ | ✓ | ✓ | ✓ | ✓ | incomplete | ✓ | ✓ | ✓ |
| AAH3920 | ✓ | ✓ | ✓ | ✓ | ✓ | ✓ | ✓ | ✓ | ✓ | ✓ | missing | missing | Missing |
| AAC2051 | ✓ | ✓ | ✓ | ✓ | ✓ | ✓ | ✓ | incomplete | incomplete | missing | ✓ | ✓ | ✓ |
| AAH1795 | missing | ✓ | ✓ | ✓ | ✓ | ✓ | ✓ | incomplete | ✓ | ✓ | incomplete | ✓ | ✓ |
| AAW0131 | missing | incomplete | ✓ | ✓ | ✓ | ✓ | ✓ | ✓ | ✓ | ✓ | ✓ | ✓ | ✓ |
| AAL9247 | incomplete | ✓ | ✓ | ✓ | ✓ | ✓ | ✓ | ✓ | ✓ | ✓ | incomplete | ✓ | ✓ |
| AAO8223 | missing | ✓ | ✓ | ✓ | ✓ | ✓ | ✓ | ✓ | ✓ | incomplete | ✓ | ✓ | ✓ |
| AAD1720 | ✓ | ✓ | ✓ | ✓ | ✓ | ✓ | ✓ | ✓ | ✓ | ✓ | ✓ | ✓ | Missing |
| AAB7912 | ✓ | ✓ | ✓ | ✓ | ✓ | ✓ | ✓ | ✓ | ✓ | ✓ | ✓ | ✓ | Missing |
| AAE6393 | incomplete | ✓ | ✓ | ✓ | ✓ | ✓ | ✓ | ✓ | ✓ | ✓ | ✓ | ✓ | ✓ |
| ABA3294 | ✓ | ✓ | ✓ | ✓ | ✓ | ✓ | ✓ | incomplete | ✓ | ✓ | ✓ | ✓ | ✓ |
| AAC2863 | incomplete | ✓ | ✓ | ✓ | ✓ | ✓ | ✓ | ✓ | ✓ | ✓ | ✓ | ✓ | ✓ |
| ABX5303 | missing | ✓ | ✓ | ✓ | ✓ | ✓ | ✓ | ✓ | ✓ | ✓ | ✓ | ✓ | ✓ |
| ABZ7255 | ✓ | ✓ | ✓ | ✓ | ✓ | ✓ | ✓ | incomplete | ✓ | ✓ | ✓ | ✓ | ✓ |
| AAA7683 | ✓ | ✓ | ✓ | ✓ | ✓ | ✓ | ✓ | ✓ | incomplete | ✓ | ✓ | ✓ | ✓ |
| AAH1501 | ✓ | ✓ | ✓ | ✓ | ✓ | ✓ | ✓ | ✓ | ✓ | ✓ | incomplete | ✓ | ✓ |
| AAD4703 | incomplete | ✓ | ✓ | ✓ | ✓ | ✓ | ✓ | ✓ | ✓ | ✓ | ✓ | ✓ | ✓ |
| AAM6304 | missing | ✓ | ✓ | ✓ | ✓ | ✓ | ✓ | ✓ | ✓ | ✓ | ✓ | ✓ | ✓ |
| ACP4019 | ✓ | ✓ | ✓ | ✓ | ✓ | ✓ | ✓ | ✓ | ✓ | incomplete | ✓ | ✓ | ✓ |
| AAZ0832 | missing | ✓ | ✓ | ✓ | ✓ | ✓ | ✓ | ✓ | ✓ | ✓ | ✓ | ✓ | ✓ |
| AAD8974 | ✓ | ✓ | ✓ | ✓ | ✓ | ✓ | ✓ | ✓ | ✓ | ✓ | incomplete | ✓ | ✓ |
| AAU9767 | missing | ✓ | ✓ | ✓ | ✓ | ✓ | ✓ | ✓ | ✓ | ✓ | ✓ | ✓ | ✓ |
| AAC1834 | ✓ | ✓ | ✓ | ✓ | ✓ | ✓ | ✓ | ✓ | incomplete | ✓ | ✓ | ✓ | ✓ |
| AAM9110 | ✓ | ✓ | ✓ | ✓ | ✓ | ✓ | ✓ | ✓ | ✓ | ✓ | ✓ | ✓ | incomplete |
| ACE9213 | missing | ✓ | ✓ | ✓ | ✓ | ✓ | ✓ | ✓ | ✓ | ✓ | ✓ | ✓ | ✓ |

### Text S2. Constructing, standardizing, and sequencing mock communities

To estimate the accuracy of SPIKEPIPE, we constructed two kinds of mock communities using known input DNA amounts of Zackenberg species: ‘mock-even’ and ‘mock-gradient’ communities (Table S2). For the six mock-even communities, we used 20 species (19 Diptera, one spider), with the same amount of DNA added from each species. Two mock-evens used 50 ng DNA per species, two used 100 ng, and two used 200 ng (Table S2). For the mock-gradient communities, we made three replicate communities, each with 19 input species (all Diptera), starting from 20 ng of DNA and increasing geometrically by a factor of 1.3 so that for the most abundant species we used 2698.9 ng of DNA, a 135-fold range of DNA biomass (Table S2). The mock-gradient mimics the common situation where some very abundant species mix with a few rare species.

#### S.2.1. Community contents

For each of the 20 species in the mock soups (Table S2), we individually extracted DNA using the same non-destructive procedure that we used for the Zackenberg environmental samples, soaking the bodies individually in lysis buffer and then purifying and extracting DNA from the soaking solution. We used this approach to be consistent with the treatment of ecological samples from Zackenberg, as described in Text S4.1. We first purified and extracted DNA from a 100 µl aliquot of soaking solution of each species. The DNA amount was quantified from these extractions using a Qubit 2.0 fluorometer (Life Technologies, USA). Each mock community was then created by pooling soaking solutions according to the quantities in Table S2, after which we aliquoted out 1.2 ml of lysis buffer from each mock community, spiked each aliquot with a fixed amount (0.2, 0.4 and 0.8 ng, respectively) of the three species included in the internal-standard DNA (see below), purified and extracted DNA from each aliquot, and sent the extractions for library-prep and sequencing. We also created and sequenced two negative-control samples containing only internal-standard DNA.

#### S.2.2. Spike-in DNA

As internal standards (see *Step 2* in the main paper), we spiked our samples with fixed quantities of DNA from three different taxa, none of which occurs in the Zackenberg area. The first one, *Bombyx mori* (Lepidoptera: Bombycidae), was bought from a food market in Kunming, China. The other two species, an unknown beetle species in Coleoptera: Mordellidae and another in Coleoptera: Elateridae, were provided by Southern China DNA Barcoding Center, Kunming, China. From each of these taxa, genomic DNA was extracted using Qiagen DNeasy Blood and Tissue Kit. PCR reactions were performed with the following protocol in a total volume of 15 μl: 4.6 μl of distilled water, 4.6 μl of 2 x MyTaq Red Mix polymerase (Bioline), 0.45 μl each of 10 μM primer (LCO1490 and HCO2198 (Folmer *et al*. 1994)) and 2 μl of the DNA template. The PCR cycling conditions were as follows: 5 min initial denaturation at 94°C, followed by 34 cycles of a 30 s denaturation at 94°C, a 30 s annealing at 52°C and a 60 s elongation at 72°C, ending with 15 min final elongation at 72°C. The PCR products were purified using Qiagen’s QIAquick PCR Purification Kit and quantified using a Qubit 2.0 fluorometer (Life Technologies, USA).

#### S.2.3. Sequencing

Low Input Transposase-Enabled (LITE) libraries were constructed from the mock-community samples and sequenced at 125 PE (250-500 bp insert size) on an Illumina HiSeq2500 at Earlham Institute (formerly TGAC) in Norwich, UK. The six mock-even communities were sequenced twice, in separate runs, allowing us to test for differences across sequencing runs. One of the even-50 mocks failed in the first sequencing run, leaving us with 11 (=5+6) total mock-even samples. The three mock-gradient communities were sequenced once.

**Table S2.**
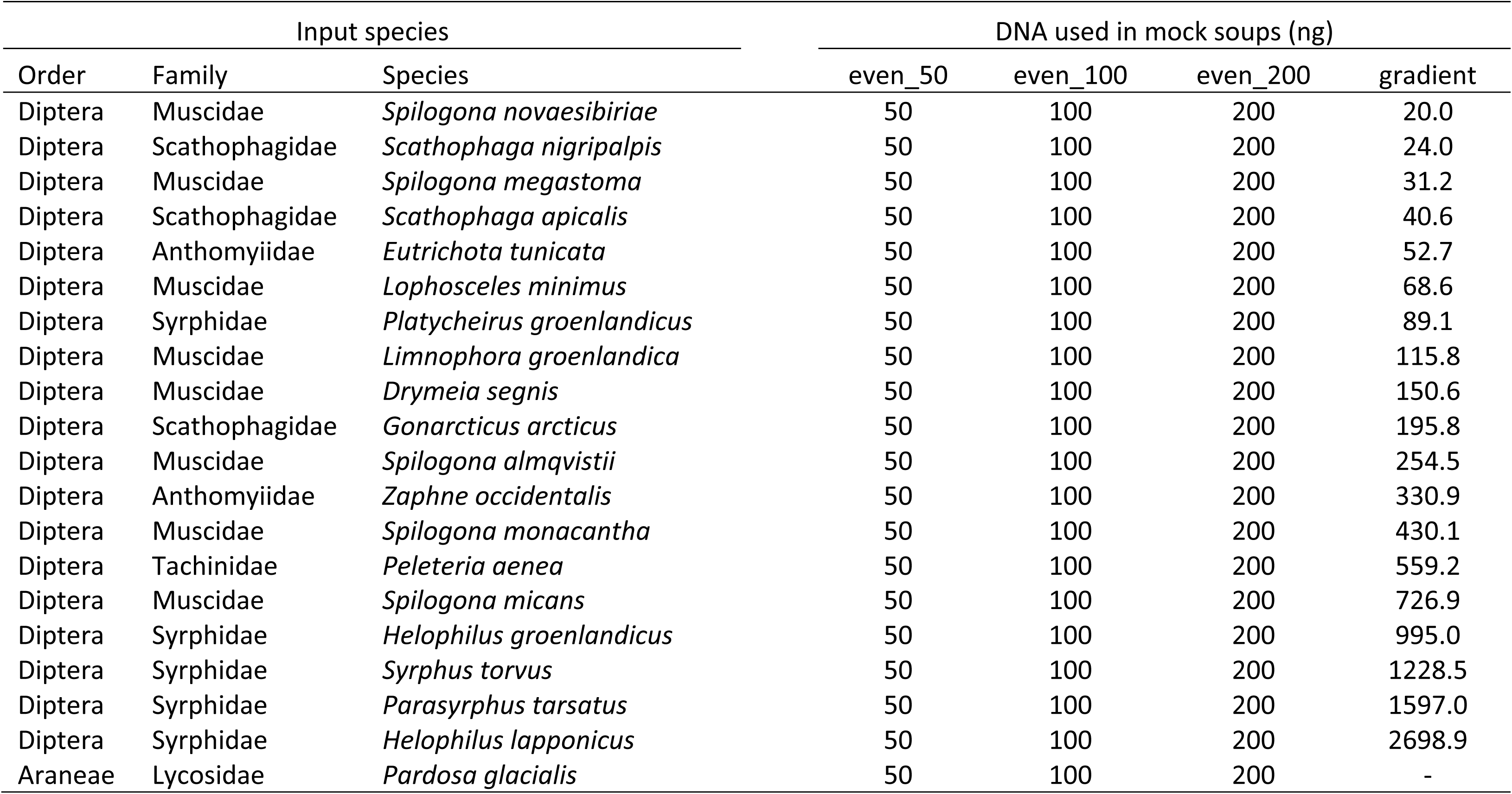
List of species and amount of input DNA used for each mock community. For the six mock-even communities, we used 20 species (19 Diptera, one spider), with DNA from each species added in the same amount. The six mock-even communities were sequenced twice (RUN EF and RUN GH). In RUN EF, one of the mock-evens (even_50 ng) failed, leaving us with 11 total mock-even datasets. For the mock-gradient communities, we made three replicate communities, each with 19 input species (all Diptera), starting from 20 ng and increasing geometrically by a factor of 1.3.

| Input species |  |  | DNA used in mock soups (ng) |  |  |  |
| --- | --- | --- | --- | --- | --- | --- |
| Order | Family | Species | even_50 | even_100 | even_200 | gradient |
| Diptera | Muscidae | *Spilogona novaesibiriae* | 50 | 100 | 200 | 20.0 |
| Diptera | Scathophagidae | *Scathophaga nigripalpis* | 50 | 100 | 200 | 24.0 |
| Diptera | Muscidae | *Spilogona megastoma* | 50 | 100 | 200 | 31.2 |
| Diptera | Scathophagidae | *Scathophaga apicalis* | 50 | 100 | 200 | 40.6 |
| Diptera | Anthomyiidae | *Eutrichota tunicata* | 50 | 100 | 200 | 52.7 |
| Diptera | Muscidae | *Lophosceles minimus* | 50 | 100 | 200 | 68.6 |
| Diptera | Syrphidae | *Platycheirus groenlandicus* | 50 | 100 | 200 | 89.1 |
| Diptera | Muscidae | *Limnophora groenlandica* | 50 | 100 | 200 | 115.8 |
| Diptera | Muscidae | *Drymeia segnis* | 50 | 100 | 200 | 150.6 |
| Diptera | Scathophagidae | *Gonarticus arcticus* | 50 | 100 | 200 | 195.8 |
| Diptera | Muscidae | *Spilogona almqvistii* | 50 | 100 | 200 | 254.5 |
| Diptera | Anthomyiidae | *Zaphne occidentalis* | 50 | 100 | 200 | 330.9 |
| Diptera | Muscidae | *Spilogona monacantha* | 50 | 100 | 200 | 430.1 |
| Diptera | Tachinidae | *Peleteria aenea* | 50 | 100 | 200 | 559.2 |
| Diptera | Muscidae | *Spilogona micans* | 50 | 100 | 200 | 726.9 |
| Diptera | Syrphidae | *Helophilus groenlandicus* | 50 | 100 | 200 | 995.0 |
| Diptera | Syrphidae | *Syrphus torvus* | 50 | 100 | 200 | 1228.5 |
| Diptera | Syrphidae | *Parasyrphus tarsatus* | 50 | 100 | 200 | 1597.0 |
| Diptera | Syrphidae | *Helophilus lapponicus* | 50 | 100 | 200 | 2698.9 |
| Araneae | Lycosidae | *Pardosa glacialis* | 50 | 100 | 200 | - |

**Figure S1.**
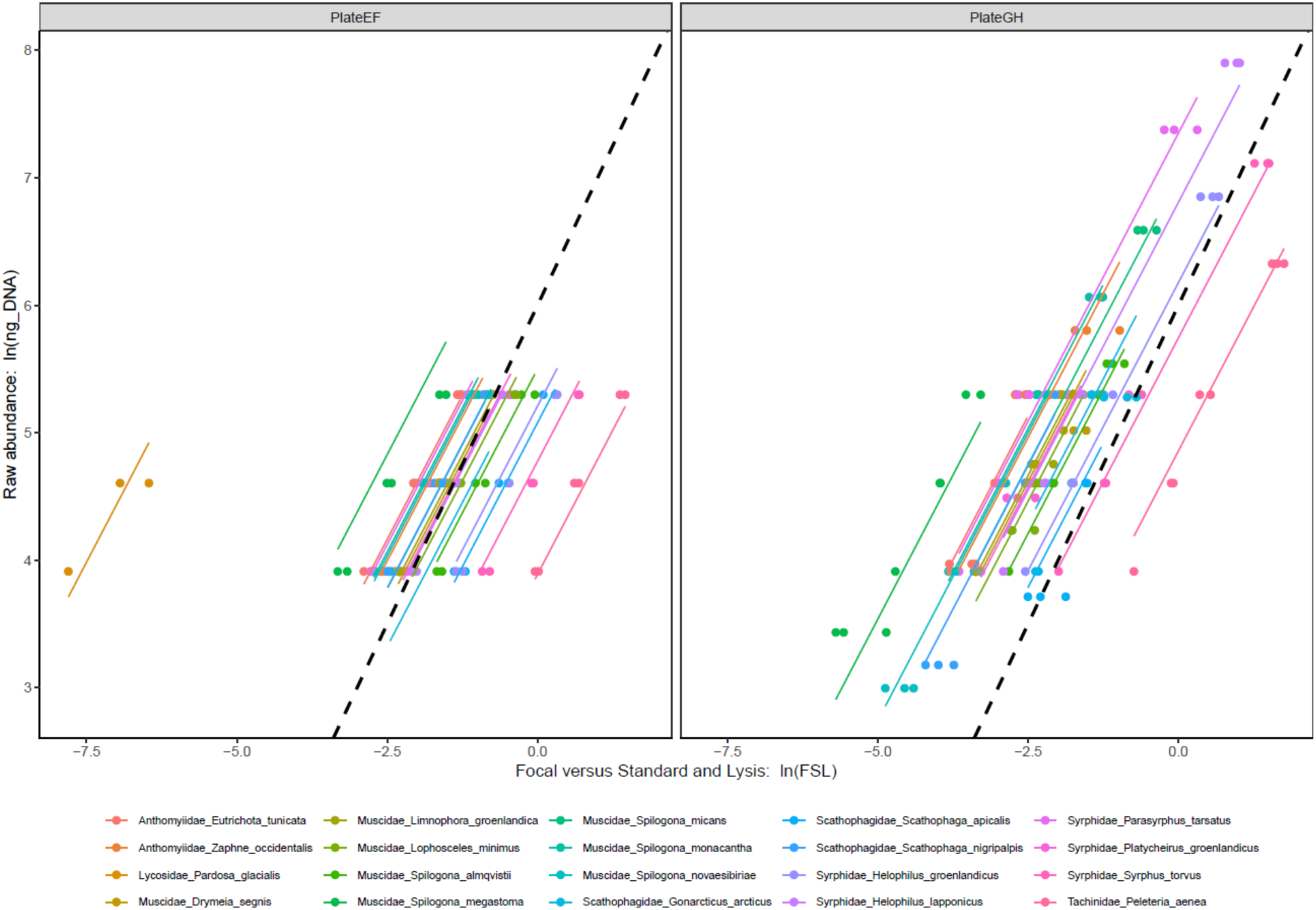
Explaining within-species variation in species abundance in the mock-community experiments. The number of mapped reads after correction for the internal standard and the fraction of lysis buffer used (FSL) predicts within-species variation in input DNA (ng). PlateEF represents the sequencing run with mock-even communities only, and PlateGH represents the run with both the mock-even and mock-gradient communities. The reference line (black dashed) in both panels has slope 1 and intercept 6 and shows that the relationship is nearly directly proportional and that there is a RUN effect. The final accepted model thus included FSL, SPECIES, and RUN (Table 2). The farthest-left species in the PlateEF run is the Arctic wolf spider, *Pardosa glacialis*.

### Text S3. Environmental samples

Arthropod samples were collected as part of the BioBasis monitoring program (Schmidt *et al*. 2016; Christensen *et al*. 2017) at the Zackenberg Research Station, located in the High-Arctic zone of northeast Greenland (74°28’ N; 20°34’ W). Arthropods were monitored weekly from 1996 to 2013 (most samples from 2010 were unfortunately lost while being transported from Greenland).

#### S.3.1. Collection protocol and storage

Altogether, arthropods were collected from five sampling plots, each plot containing eight pitfall traps during 1996–2006 (A, B, C, D, E) and four traps during 2007–2013 (A, B, C, D). All traps were emptied on a weekly basis throughout the summer season of each year. When the traps were emptied, the trap liquid was poured through a small aquarium net into a spare cup, whereupon the liquid was poured back into the repositioned upper cup. The catch was then emptied into a 10-cl vial with alcohol by turning the net inside-out inside the vial. All remaining invertebrates were carefully removed from the aquarium net with tweezers and moved to the container. After each sampling day, the net was rinsed in fresh water, and at the beginning of each sampling season (year), the net was exchanged for a new one.

Arthropods were sorted and counted by technicians from the Department of Bioscience at Aarhus University, Denmark. At this stage, the arthropods were resolved not to species level, but to rough taxonomic groups spanning species of different ecologies (Schmidt *et al*. 2016). (For exact taxonomic groups, see Table S3.). All specimens were subsequently stored in 75% ethanol in room temperature at the Museum of Natural History, Aarhus. Weekly samples were stored in vials closed with cotton in larger 300-500ml jars filled with ethanol (see Fig. S2). Importantly, the larger jar contained multiple tubes of the taxonomic fraction in question (see Table S3, Fig. S2).

We note that the processing and storage of samples – as is typical of samples collected for monitoring and later stored in wet collections – contain several sources of contamination to control for. First, the use of a single set of equipment for sieving the samples in the field allow fragments of arthropods to be mixed between samples. Second, the storage of samples in open tubes in shared vials filled with ethanol (Fig. S2) probably allows some mixing of DNA, since the sample fluid has been shown to contain DNA of samples (Hajibabaei *et al*. 2012). This problem is compounded by the renewal of ethanol to keep the medium “fresh”, as undertaken every year.

From this larger monitoring design (Schmidt *et al*. 2016), we used for this study arthropods collected weekly throughout the summer from 1997 to 2013 from three individual traps (A, B, C) located in a single sampling plot “Art3”. This plot is located in mesic heath dominated by lichens (an almost complete cover of organic crust) and white Arctic bell-heather *Cassiope tetragona*, and with scattered individuals of Arctic willow *Salix arctica* and Arctic blueberry *Vaccinium uligonosum* (Schmidt *et al*. 2016). We used data from samples collected from 1997 to 2013. (Samples from 1996 were omitted as the samples from this first year of the biomonitoring program had been somewhat differently handled, sorted, and labelled.) The resulting set of samples consisted of an estimated 23,001 arthropod individuals. We omitted Collembola, Acari, and all larval stages from our analyses, since these taxa are not reliably sampled by yellow pitfalls, nor had they been consistently sorted by technicians (being sometimes stored, sometimes not).

**Figure S2.**
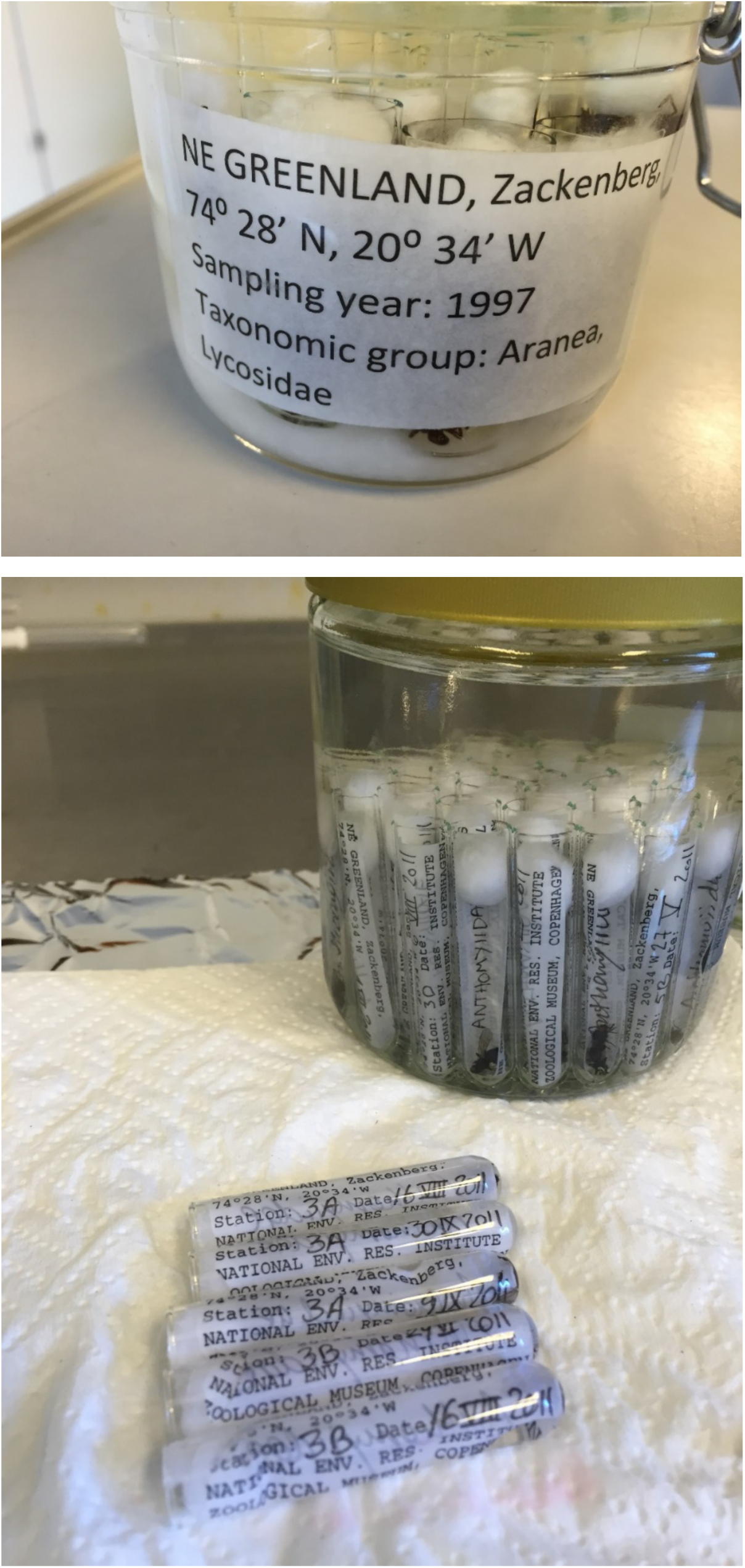
Sample storage. Samples of the compound taxonomic fractions identified in Table S3 were stored in tubes in shared vials filled with ethanol. Each tube was closed by a wad of cotton. Note that the collection includes a range of tube (ca. 3 ml to 15 ml) and vial sizes, with a few representative examples shown here.

**Table S3.**
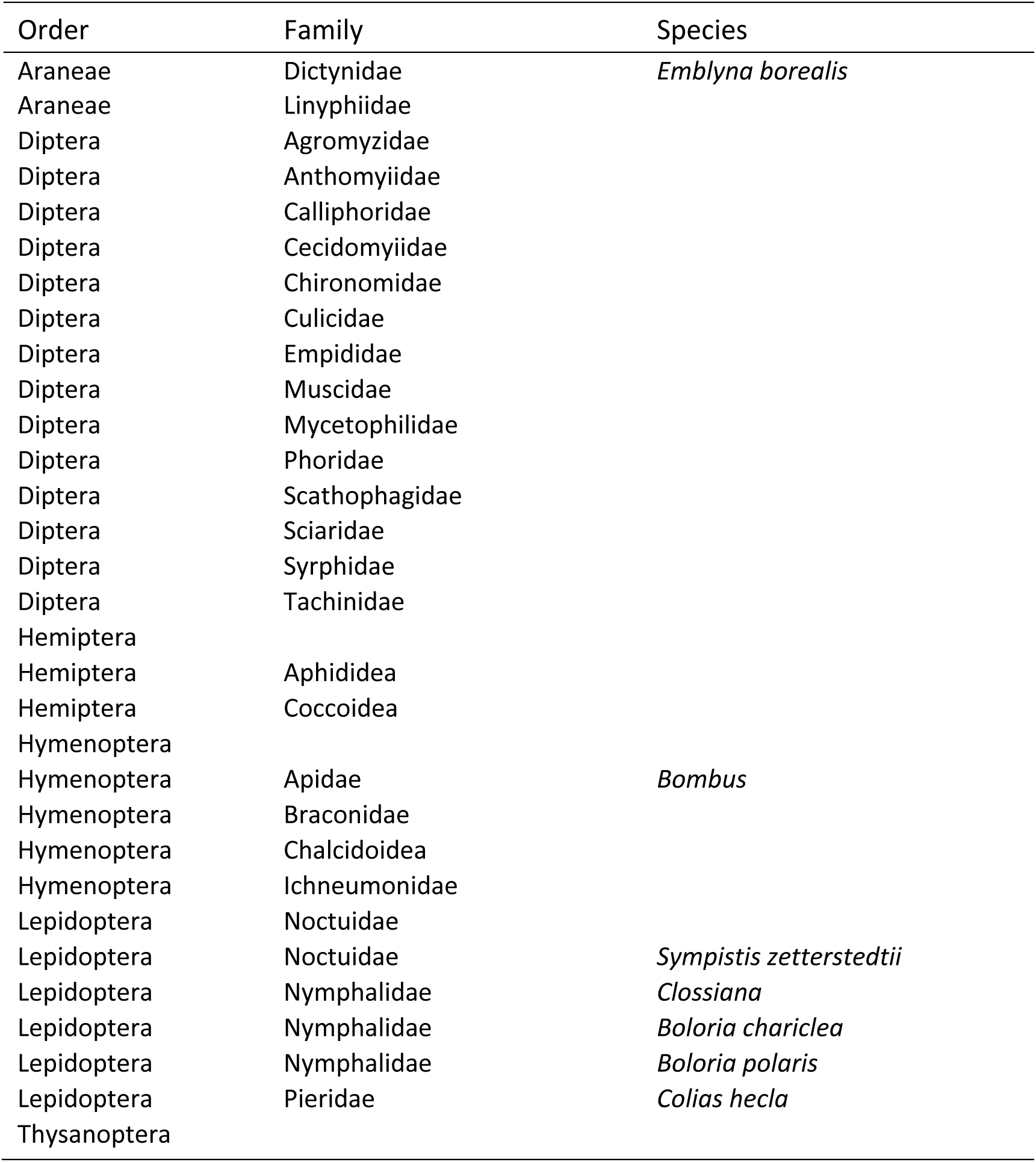
Taxonomic classification used in the Zackenberg collection. The table identifies the taxonomic ranks to which samples are sorted, following Schmidt *et al*. (2016). Note that most fractions correspond to families but that in some cases, only a single species of this family will occur in the region, thus resulting in de facto species-level identification. For a full list of species encountered at Zackenberg, see Wirta *et al*. (2015). Note that the table includes only taxonomic fractions encountered in the subset of samples processed by us; for the full set of taxonomic fractions used in the morphology-based sorting of specimens being part of the biomonitoring programme, see Schmidt *et al*. (2016).

### Text S4. Processing of bulk samples from the Zackenberg collection

The Zackenberg collection is, to our knowledge, the only systematically curated, long-term data set of arthropod communities from the High Arctic. Thus, we designed our protocol to retain the original structure of the full collection, with all individuals left as intact as possible from a morphological perspective. Both DNA extracts and full sequencing output are archived for future studies.

#### S.4.1. DNA extraction

Since the original Zackenberg pitfall-trap samples were sorted to taxonomic fractions (see above and Table S3), our first task was to re-pool whole trap-week samples from the separate trap-week-taxon sub-samples (Table S3). For each trap-week-taxon sub-sample test tube, we did digestion, the first step of DNA extraction, separately. Before digestion, we counted and air-dried the contents of each tube on qualitative filter paper (Whatman, USA). We then used a non-destructive method of DNA extraction: after drying, we soaked all individuals in lysis buffer (modified from Gilbert *et al*. 2007 and consisting of 3 mM CaCl_2_, 2% Sodium Dodecyl Sulfate, 40 mM Dithiothreitol, 250 ug/ml proteinase K, 40 mM Tris buffer pH 8 and 100 mM NaCl) corresponding to 5 times the volume of individuals in each tube and incubated the specimens for 72 hours in a waterbath shaker rotating at 200 RPM with the temperature set to 56 °C. Subsequently, we returned the exoskeletons specimens to their original ethanol-filled vials in the collection, thereby restoring its original organization.

To recreate the original trap-week samples for downstream analyses, we pooled the lysis buffers of all the tubes from the same week and same trap, and we recorded the total volume of lysis buffer. After pooling, we aliquoted 1200 μl of lysis buffer from each pooled sample for purification. For samples with less than 1200 μl of lysis buffer, we used the total volume for purification. Before purification, we added a standard spike-in of three species not found in the Zackenberg fauna to serve as an internal standard (see *Step 2* of the main paper, with details on how these spike-ins were prepared in Text S2.2 above). The resulting spiked, trap-week aliquots were purified using Qiagen’s QIAquick PCR Purification Kit.

Samples from the earliest and latest parts of the arctic summer were often very small, comprising a few individuals only. For these, we individually DNA-barcoded each individual with PCR primers LCO1490 and HCO2198 (Folmer *et al*. 1994) and identified to species by comparing the Sanger sequences to the BOLD database (The Barcode of Life Data Systems, www.barcodinglife.org, Ratnasingham & Hebert 2007) – which, due to the work of Wirta *et al*. (2015), includes DNA barcodes of nearly all arthropod species present in the target region. This supplementary count dataset will be added to the final dataset used for ecological analyses and is omitted from all analyses in this paper, as this paper focused on the mitogenome pipeline.

#### S.4.2. Library prep and Illumina Sequencing

A separate Low Input Transposase-Enabled (LITE) sequencing library was constructed for each of the 493 trap-week samples at the Earlham Institute (formerly TGAC) in Norwich, UK. LITE libraries are low-cost (£5 each in 2017), are produced in an automated pipeline in standard 96-well plates, and are thus purchased 96 libraries at a time. Over 2016 and 2017, we altogether sequenced 712 trap-week samples, submitted to EI as three pairs of 96-well plates (Table S4.2.1).

- Plates A and B contained trap-week samples from 1997-1999 and 2011-2013.
- Plates E and F contained samples from 2000-2002 and 2007-2009, plus one sample from 1998, one sample from 1999, 4 trap-week technical replicates from 1998, 1999, and 2013, and the first batch of mock-evens.
- Plates G and H contained samples from 2003-2006, 4 or 5 trap-week technical replicates from the other previous years, plus the second batch of mock-evens, the mock-gradients, and two negative controls.

We used the replicates to correct for stochasticity in sequencing depth across sequencing runs. Each pair of plates was sequenced over 12 lanes of an Illumina HiSeq 2500 at 125 bp PE with a 250-500 bp insert size. After determining that the internal-standard spike-in took up a large proportion of the dataset in Plates A and B (see main text), we resequenced them on another 11 total lanes to augment the dataset (Plates A2 and B2), but with some missing samples because their original DNA was no longer sufficient to construct new libraries.

The two negative controls in Plates G and H were used to test for the unlikely possibility that the LITE library preparation pipeline causes sample cross-contamination. As expected, the two negative controls resulted in no mapped reads, except to the internal-standard species. Note that amplicon-sequencing protocols require extensive use of negative controls, because trace DNA contamination during PCR set up can result in large numbers of artefactual reads, but this is not an issue for shotgun-sequencing protocols like ours.

**Table S4.2.1.**
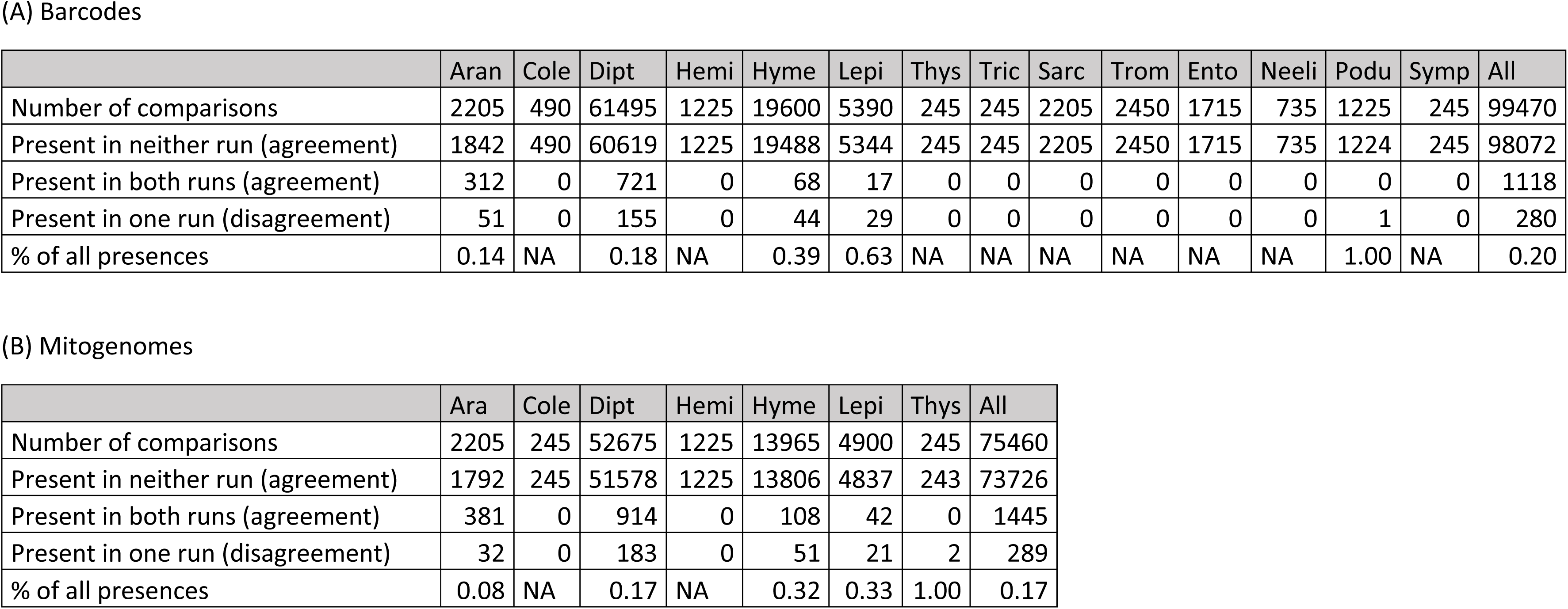
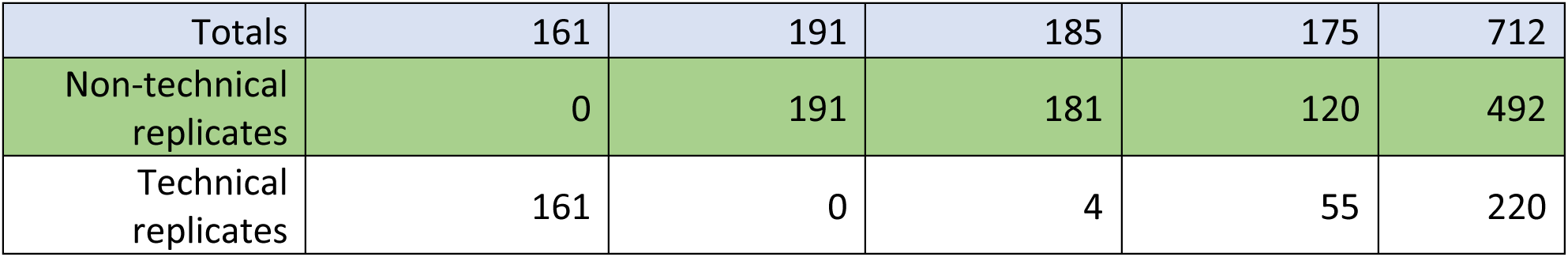
Distribution of trap-week samples by year and sequencing run. Each pair of plates was submitted separately for a sequencing run (e.g. Plates A & B containing years 1997-1999 and 2011-2013 were sequenced in “Run AB”). Green cells indicate the original trap-week samples. White cells indicate technical replicates (thus, RunA2B2 contains a nearly complete set of technical replicates of Run AB). Two samples (light green cells) were successfully sequenced later than their originally designated run (e.g. a 1999 sample was sequenced in Run EF, after failing in Run AB). The total number of trap-week samples sequenced at least once (unique environmental samples) is 492.

#### S.4.3. Taxonomic impacts on performance

The Zackenberg samples are dominated by Diptera, which is why our mock samples included a high content of such taxa. In validating the repeatability of results across replicate runs of environmental samples, we repeated the analyses of Table 3 of the main paper separately for all orders present in our data. The results (Table S4.3.1) show that Diptera and Araneae are the most common orders, and that the consistency among replicate samples is somewhat higher for Araneae than for Diptera. For Hymenoptera and Lepidoptera, the consistency is somewhat lower than for Diptera, but these orders are too rare to make robust conclusion.

**Table S4.3.1.**
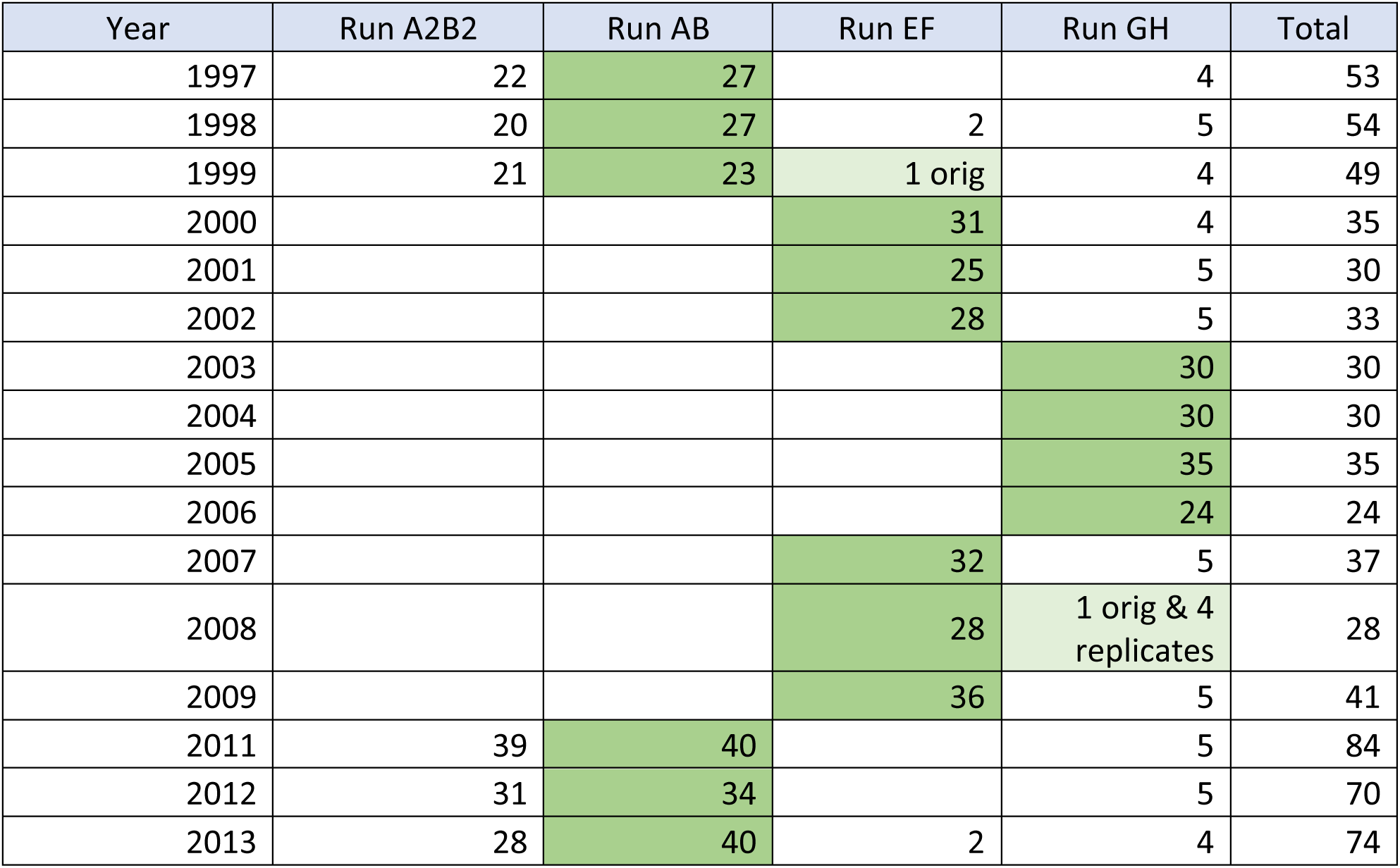
Consistency of re-sequenced environmental data across arthropod orders. This table recreates the results of Table 3 in the main text, but subdivided into different orders, for mapping (a) to DNA barcodes and (b) to mitogenomes. In other words, we show the consistency in species presences and absences between two sequencing runs, with the total number of comparisons equaling the total number of sample-by-species combinations. To evaluate the robustness of detecting species presence, we counted how often a species was inferred to be present or absent in both cases (agreement) or present in one case only (disagreement). The proportion of presences is counted as the fraction of all samples where the species was inferred to be present in at least one run. Abbreviations: Aran=Araneae, Cole=Coleoptera, Dipt=Diptera, Hemi=Hemiptera, Hyme=Hymenoptera, Lepi=Lepidoptera, Thys=Thysanoptera, Tric=Trichoptera, Sarc=Sarcoptiformes, Trom=Trombidiformes, Ento=Entomobryomorpha, Neeli=Neelipleona, Podu=Poduromorpha, Symp=Symphypleona.

### Text S5. Mapping reads against references

We removed remaining Illumina adapter sequences using *TrimGalore* 0.4.5 (--paired --length 100 -trim-n) (Martin 2011) and used *minimap2* 2.10 in short-read mode (-ax sr) (Li 2018) to map the paired-end reads from each sample against a fasta file containing the 308 mitogenomes and the 3 internal-standard barcode sequences. We used *samtools* 1.5 (Li *et al*. 2009) to sort, convert to bam format, exclude reads that were unmapped or mapped as secondary alignments and supplementary alignments, and include only ‘proper-pair’ read mappings (mapped in the correct orientation and at approximately the correct distance apart) at ≥ 48 ‘mapping quality’ (*MAPQ*) (view sort -b -F 2308 -f 0×2 -q 48), where *MAPQ* = −10log_10_(prob that mapping position is wrong). We accepted *MAPQ* ≥ 48 after inspection of the highly bimodal distribution of quality values, with most reads giving *MAPQ* = 0 (i.e. maps well to multiple locations) or 60 (probability of error = 0.0001%). *MAPQ* = 48 corresponds to a probability of error equaling ca. 0.001%. Informally, we found that limiting quality to only the highest value, 60, had little effect on the results, whereas including low-quality mappings (-q 1) led to more false-positive hits (data not shown).

The output for each sample is the number of mapped reads per mitogenome and internal-standards sequences that have passed the above filters. However, it is still possible for a mitogenome to receive false-positive mappings. Thus, we add another round of filtering. We expect that if a species is truly in a sample, mitochondrial reads from that sample will map along the length of that species’ mitogenome, resulting in a higher percentage coverage. In contrast, if reads map to just one location on a mitogenome, even at high *MAPQ*, the percentage coverage will be low, and we consider those mappings to be false-positive detections caused by that mapped portion of the mitogenome being very similar to a species that is in the sample but not in the reference database (Fig. S3). We used *bedtools* 2.27.1 to calculate the number of overlapping reads at each position along the reference sequence (genomecov -d). The percentage coverage (henceforth PC) is the fraction of positions covered by one or more mapped reads.

**Figure S3.**
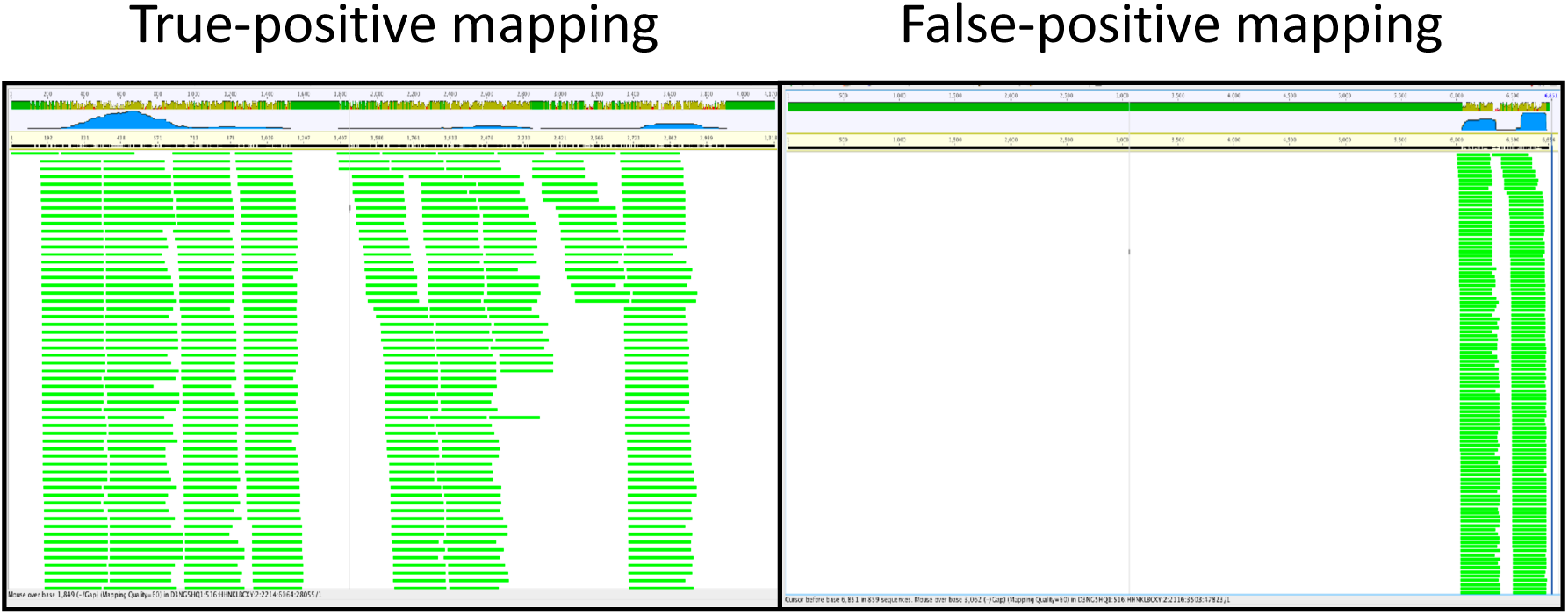
Visual example of true- and false-positive mapping of reads to mitogenomes. In the false-positive example, there are many reads, but they together only cover a small percentage of the total mitogenome length, indicating that this portion of the mitogenome sequence is probably conserved with another species that is not present in the mitogenome reference database.

### Text S6. How to run Step 4 (bioinformatics) of the SPIKEPIPE pipeline

Here we describe how to the user may apply SPIKEPIPE pipeline to a subset of the data used in this paper, henceforth called ArcDyn tutorial.

To install the ArcDyn tutorial, download and untar the 4.9 GB tutorial file (`ArcDyn_tutorial_20190126.tar.gz`) to the root directory by running this on the command line:

`cd ∼/`

`curl -O` https://s3.eu-west-2.amazonaws.com/arcdyntutorial20190126/ArcDyn_tutorial_20190126.tar.gz

`tar -xzvf ArcDyn_tutorial_20190126.tar.gz`

After unzipping, you will have a new folder called `ArcDyn_tutorial/`.

Navigate to `ArcDyn_tutorial/_tutorial_start_here/`

#### S.6.1. Bioinformatic and Statistical scripts for ArcDyn tutorial

**Overview.** The input data for the bioinformatics scripts are found in the folders `sequences_files/` (the sequence files of the environmental samples), `reference_seqs/` (the reference mitogenomes), and `reference_files/` (the sample metadata, plus examples of the Unix scripts that were used to prepare the raw sequence files for the tutorial). The scripts are located in the folder `_tutorial_start_here/`, and they utilize the software and utilities described below. The tutorial data are a subset of the full dataset (mock samples from PlatesGH only and environmental samples from Year 2006 only), to prevent excessive file size.

**`Step_4.1_read_mapping_and_filtering.sh`** – This script runs in the Unix shell and (1) uses `seqtk` to count the total number of reads per fastq file, (2) uses `minimap2` to map the reads in each sample to the 308 mitogenomes, (3) uses `samtools` to filter the mapped reads to include only high quality mappings and count the number of mapped reads to each mitogenome in each sample (`*_idxstats.txt`), and (4) uses `bedtools` to count sequencing depth for each position on each mitogenome in each sample (`*_genomecov_d.txt.gz`)

**`Step_4.2_read_in_samtools_output_and_metadata.Rmd`** – This script reads in the `*_idxstats.txt` and `*_genomecov_d.txt.gz` files, calculates percent coverages of the mappings, combines them with the sample metadata (e.g. date, trap, sequencing run, lysis buffer proportion), does sanity checks, removes any failed samples, and separates the environmental samples from the mock samples. The output is a single table of all environmental samples and a single table of all mock samples: `idx_meta_genomecov_PlatesGH_20190115_308mitogenomes.txt` and `mocks_idx_meta_genomecov_PlatesGH_20190125_308mitogenomes.txt`. I include a date (`YYYYMMDD`) in the filename to indicate when the read-mapping run occurred (here, `20190125`), and I indicate the reference sequences used (here, `308_mitogenomes`).

**`Step_4.3_format_datafiles_for_Step5_statistical_analysis.Rmd`** – This script runs in *R* and reads in the two output files from **`Step_4.2`**, plus the `fastq_read_counts_ArcDyn_tutorial.txt file`, and generates the multiple data files used in the statistical analysis, saving them in a single RData file: `input_data_step5_subset.RData.`

**`Step_5.1_mock_analysis.r`** – This script runs in *R* and reads in `input_data_step5_subset.RData`. The script fits statistical models predicting species occurrences and abundances from the mock data, and generates outputs analogous to Tables 1 and 2 in the main text. Because the tutorial dataset contains only one year of environmental data, the tutorial does not run **`Step_5.2_environmental_analysis.r`**, which requires multiple years of data. The environmental analysis can be run on the full dataset, which is described in S7 below.

#### S.6.2. Software installation for ArcDyn tutorial

**Software installation.** Here we describe software packages used in the pipeline (not including the R packages, which are listed in the R scripts (`2_idxstats_tabulate_macOS_PlatesGH.Rmd, 3_format_datafiles_for_statistical_analysis.Rmd`). If you are installing with Homebrew on macOS, the path to each package is set automatically, and the software can be called without its pathname. If you install manually, the path can be set before running, allowing the software to be called without its preceding pathname. For instance, assuming that the `minimap2` binary is installed on your computer at `∼/src/minimap2/`, run this in your script:

`PATH=$PATH:∼/src/minimap2/`

Many Unix packages can be installed on macOS using Homebrew (https://brew.sh)

**s3cmdtools**: for managing Amazon S3 from the command

https://s3tools.org/s3cmd

**sed**: stream editing utility in Unix. Note that sed works differently on macOS and Linux. The way to get the Linux version of sed is to install GNU sed on macOS: `gsed`. On macOS, install with homebrew: `brew install gnu-sed`

**GNU parallel**: This utility allows a single command to generate multiple jobs running in parallel, each one with different inputs (and/or command options). It is an alternative to file-globbing wildcards and loops.

https://www.gnu.org/software/parallel/

On macOS, install with homebrew: `brew install parallel`

**TrimGalore**: This is a wrapper for cutadapt and optionally, fastqc

https://www.bioinformatics.babraham.ac.uk/projects/trim_galore/

https://github.com/FelixKrueger/TrimGalore/archive/0.5.0.zip

**cutadapt**: This package trims away Illumina adapters and filters read pairs by length and quality

https://github.com/marcelm/cutadapt/

On macOS, install with homebrew: `brew install brewsci/bio/cutadapt`

**fastQC**: This package summarises quality and data size statistics for fastq files

https://www.bioinformatics.babraham.ac.uk/projects/fastqc/

On macOS, install with homebrew: `brew install fastqc`

**multiqc**: This package summarises the outputs of multiple genomics packages. We use it to summarise the fastQC results installation instructions on: https://multiqc.info

**minimap2**: We use this package for mapping reads to mitogenomes, instead of the more commonly used bwa. Both minimap2 and bwa are written by the same author, Li Heng, but according to Li Heng: “For >100bp Illumina short reads, minimap2 is three times as fast as BWA-MEM and Bowtie2, and as accurate on simulated data.”

https://github.com/lh3/minimap2

On macOS, install with homebrew (or place the binary from github in a convenient location and set the path, since homebrew currently throws an error with this one): `brew install brewsci/bio/minimap2`

**samtools**: We use this package to filter and sort bam files and to generate a bam index file. http://www.htslib.org

https://github.com/samtools/samtools/releases

On macOS, install with homebrew: `brew install samtools`

**bedtools**: This is a set of utilities for managing and extracting information from bed and bam files.

https://bedtools.readthedocs.io/en/latest/

https://bedtools.readthedocs.io/en/latest/content/installation.html

**R**: The standard software for statistical language and a programming environment.

https://cran.r-project.org

**RStudio**: A GUI for R.

https://www.rstudio.com

**Utilities.** SPIKEPIPE uses these two packages:

**seqkit**: Creates a nicely formatted table of data sizes and read numbers per fastq file. https://github.com/shenwei356/seqkit

On macOS, install with homebrew: `brew install brewsci/bio/seqkit`

**seqtk**: Generally useful fastq/fasta file utility set. I used this to downsample the fastq files.

https://github.com/lh3/seqtk

On macOS, install with homebrew: `brew install seqtk`

### Text S7. How to run Step 5 (statistics) of the SPIKEPIPE pipeline

**Overview.** The scripts and full dataset for the statistical part of SPIKEPIPE are included in Supplementary Information.

**`input_data_step5.RData`** – This data file was generated by applying the full bioinformatic scripts (github.com/dougwyu/ArcDyn) to all mock and environmental samples (1997-2013). The output of the two statistical scripts below are the analyses presented in the paper in Tables 1-3 and Figures 1-2.

**`Step_5.1_mock_analysis.r`** – This script runs in *R* and reads in `input_data_step5.RData`. The script fits statistical models predicting species occurrences and abundances to the mock data, and generates the results of Tables 1 and 2 as output.

**`Step_5.2_environmental_analysis.r`** – This script runs in *R* and applies the fitted models from the mock data used in `Step_5.1` to the environmental data and produces as output the results shown in Table 3 and Figures 1 and 2. For more details, see the comments included in the scripts.

